# Myoscaffolds reveal laminin scarring is detrimental for stem cell function while sarcospan induces compensatory fibrosis

**DOI:** 10.1101/2022.07.07.497559

**Authors:** Kristen M. Stearns-Reider, Michael R. Hicks, Katherine G. Hammond, Joseph C. Reynolds, Alok Maity, Yerbol Z. Kurmangaliyev, Jesse Chin, Adam Stieg, Nicholas A. Geisse, Sophia Hohlbauch, Stefan Kaemmer, Lauren R. Schmitt, Thanh T. Pham, Ken Yamauchi, Bennett G. Novitch, Roy Wollman, Kirk C. Hansen, April D. Pyle, Rachelle H. Crosbie

**Affiliations:** Department of Integrative Biology and Physiology, University of California, Los Angeles, Los Angeles, CA 90095, USA; Department of Microbiology, Immunology, and Molecular Genetics, David Geffen School of Medicine, University of California, Los Angeles, Los Angeles, CA 90095, USA; Eli and Edythe Broad Center of Regenerative Medicine and Stem Cell Research, University of California, Los Angeles, Los Angeles, CA 90095, USA; Department of Chemistry and Biochemistry, University of California, Los Angeles, Los Angeles, CA 90095, USA; Institute for Quantitative and Computational Biology, University of California, Los Angeles, Los Angeles, CA 90095, USA; Department of Biological Chemistry, HHMI, David Geffen School of Medicine, University of California, Los Angeles, Los Angeles, CA 90095, USA; California NanoSystems Institute, University of California, Los Angeles, Los Angeles, CA 90095, USA; NanoSurface Biomedical, Seattle, WA 98195, USA; Asylum Research, An Oxford Instruments Company, Santa Barbara, CA, 93117, USA; Park Systems, 3040 Olcott St, Santa Clara, CA 95054, USA; Department of Biochemistry and Molecular Genetics, University of Colorado, Denver, Aurora, CO 80045, USA; Department of Neurobiology, David Geffen School of Medicine, University of California, Los Angeles, Los Angeles, CA 90095, USA; Molecular Biology Institute, University of California, Los Angeles, Los Angeles, CA 90095, USA; Intellectual and Developmental Disabilities Research Center, David Geffen School of Medicine, University of California Los Angeles, Los Angeles, CA 90095, USA; Department of Neurology, David Geffen School of Medicine, University of California Los Angeles, Los Angeles, CA 90095, USA

**Keywords:** collagen, Duchenne muscular dystrophy, extracellular matrix, human pluripotent stem cells, integrin, laminin, myoscaffold, regeneration, sarcospan, skeletal muscle progenitor cells, stem cells.

## Abstract

We developed an on-slide decellularization approach to generate acellular extracellular matrix (ECM) scaffolds that can be repopulated with various cell types to interrogate cell-ECM interactions. Using this platform, we investigated whether fibrotic ECM scarring affected human skeletal muscle progenitor cell (SMPC) functions that are essential for myoregeneration. SMPCs exhibited robust adhesion, motility, and differentiation on healthy muscle-derived myoscaffolds. All SPMC interactions with fibrotic myoscaffolds from dystrophic muscle were severely blunted including reduced motility rate and migration. Furthermore, SMPCs were unable to remodel laminin dense fibrotic scars within diseased myoscaffolds. Proteomics and structural analysis revealed that excessive collagen deposition alone is not pathological, and can be compensatory, as revealed by overexpression of sarcospan and its associated ECM receptors in dystrophic muscle. Our *in vivo* data also supported that ECM remodeling is important for SMPC engraftment and that fibrotic scars may represent one barrier to efficient cell therapy.

## INTRODUCTION

Skeletal muscle is a highly adaptive tissue with a robust capacity for regeneration due to resident muscle stem cells, or satellite cells that lie beneath the basement membrane, adjacent to myofibers ^1-3^. In healthy muscle, myofiber hypertrophy from resistance exercise occurs after myofiber damage, resulting in local inflammation, satellite cell activation, and extracellular matrix (ECM) deposition leading to increased muscle mass and force production ^4^. Duchenne muscular dystrophy (DMD), a progressive muscle wasting disease, is characterized by repeated contraction-induced injury of fragile myofibers resulting in asynchronous cycles of degeneration and regeneration ^5-7^. In early stages of DMD, the muscle is able to recover following injury and maintains its regenerative capacity. However, over time, there is progressive loss of muscle function and failed regeneration, with DMD patients exhibiting muscle weakness by the age of four and loss of ambulation by age thirteen. Over time, muscle is asymmetrically replaced by fat and connective tissue as a function of increased ECM deposition, or fibrosis. The factors that contribute to myofiber decompensation and failed regeneration are unclear; however, identification of genetic modifiers of DMD disease may provide insights. To date, SPP1 (osteopontin) ^8^ and LTBP4 (latent TGF-β binding protein 4) ^9^ have been identified as genetic modifiers in both human DMD and murine models of disease. Both proteins modulate ECM production, suggesting a key role for the ECM in disease severity and progression. Osteopontin is a secreted matricellular glycoprotein that signals partially through integrin receptors and its downregulation is associated with increased strength, reduced fibrosis, and milder muscle pathology ^10,11^. LTBP4 is a regulator of the TGF-β pathway that promotes ECM protein synthesis and suppresses the activity of matrix metalloproteinases ^12,13^. Upregulation of LTBP4 is associated with increased fibrotic tissue deposition and predicts the age at loss of ambulation ^9^. These findings reveal that ECM modifiers in the presence of muscle pathology are a determinant of disease progression; however, the repair process in injured muscle involves the coordinated activities of multiple cell types, making it difficult to determine if inherent properties of the fibrotic ECM have detrimental effects on muscle regeneration.

The relationship of the ECM with contracting myofibers is far more complex relative to other tissues. The primary contact of myofibers with the ECM occurs through laminin-binding receptors that are localized into regularly repeating structures called costameres^14^. Moreover, the costameres at the cell surface interact with the repeating Z-disc component of the intracellular sarcomere, the contractile unit in muscle, through the actin cytoskeleton. The major laminin binding receptors in muscle are α7β1 integrin and the dystrophin-glycoprotein complex, which contribute to the polymerization of laminin and assembly of the basement membrane layer within the ECM ^15,16^. Disruption of this highly organized laminin-cytoskeleton network diminishes contractile function, increases muscle susceptibility to injury, and is the major underlying cause of many muscular dystrophies. Loss of dystrophin in DMD causes laminin disorganization in the basement membrane ^17^, which may have direct effects on satellite cells that require laminin degradation to initiate myogenesis ^18^. Henry, Campbell, and colleagues showed that upregulation of adhesion complexes in dystrophic muscle restored laminin organization in the basement membrane, leading to improved myofiber adhesion and reduced damage ^19^.

The major objective of the current study was to develop a reductionist approach to investigate the effect of the skeletal muscle ECM on specific functions of stem cells that are necessary for regeneration and, reciprocally, to determine how cells interact with and modulate the ECM. We tailored *in vitro* methods ^20-23^ to develop on-slide decellularization of skeletal muscle yielding acellular ECM scaffolds, or myoscaffolds, that retain native architecture and composition. We then repopulated healthy and fibrotic myoscaffolds with human skeletal muscle progenitor cells (SMPCs) derived from human pluripotent stem cells (hPSCs). Analysis revealed ECM scarring around necrotic myofibers in *mdx* myoscaffolds that had a detrimental effect on all aspects of SMPC function and induced expression of cell stress markers, while suppressing expression of cell differentiation genes in SMPCs. We additionally identified regions of the *mdx* myoscaffolds without dense scar formation, characterized by muscle hypertrophy and minimal ECM deposition, that supported robust SMPC adhesion, proliferation, migration, and differentiation. These data reveal that there are multiple states of skeletal muscle fibrosis that possess either compensatory properties, capable of supporting effective regeneration, or pathological properties that are sufficient to negatively regulate stem cell regenerative capacity.

We next analyzed myoscaffolds generated from *mdx* muscle engineered to overexpress laminin-binding receptors (*mdx* ^TG^) that restored attachment of the myofiber to the ECM and prevented muscular dystrophy ^24,25^. Surprisingly, the *mdx* ^TG^ myoscaffolds exhibited fibrosis characterized by increased collagen deposition in the absence of pathology, supporting the conclusion that excessive ECM deposition alone is not detrimental to muscle regeneration and may be compensatory in the transgenic model. The *mdx* ^TG^ myoscaffolds supported robust SMPC function and were readily remodeled by the SMPCs. Proteomic and biochemical analysis of all myoscaffolds led to the conclusion that laminin scarring, not collagen abundance, is a primary factor limiting stem cell regenerative capacity in muscle, and this was further demonstrated by SMPC breakdown of collagen but not laminin *in vitro*. These cellular behaviors were not unique to hPSC derived SMPCs as primary mouse satellite cells exhibited similar dysfunction when cultured on *mdx* myoscaffolds. Engraftment experiments in wild-type and *mdx* mice further revealed that SMPCs readily remodel the *in vivo* microenvironment, but were unable to remodel the fibrotic scars in *mdx* mice. In conclusion, the myoscaffold platform provides an *in vitro* model for testing cell-ECM interactions that may be beneficial for interrogating the efficacy of cell-based therapies.

## RESULTS

### On-slide decellularization yields acellular myoscaffolds that recapitulate ECM architecture and composition

Decellularization of entire skeletal muscles has been employed to generate acellular ECM scaffolds ^26-28^; however, this approach has many challenges, including the disproportionate removal of cellular material due to unequal detergent exposure in the peripheral versus central regions of the tissue. To address these limitations, we developed an on-slide decellularization method in which transverse muscle cryosections were mounted on microscope slides prior to decellularization, permitting equal detergent exposure across the tissue surface. The DMD murine model (*mdx*) of chronic muscle injury permits investigation of multiple stages of disease progression from early-stage, characterized by hypertrophic myofibers and minimal fibrosis, to late-stage, characterized by myofiber necrosis and fibrotic scars, all within the same tissue ^29,30^. Murine *mdx* quadriceps muscle was selected for analysis as it has been widely studied and is a clinically relevant muscle affected in DMD ^31-33^. The effectiveness of decellularization using this method, including the extent of cellular removal and the preservation of ECM architecture, was assessed with hematoxylin and eosin (H&E) staining of sections from wild-type (WT) controls after decellularizing for 10 to 60 minutes (Fig. 1a). Complete decellularization of both WT and *mdx* samples was achieved after 30 minutes in 1% SDS, as revealed by the absence of myofibers and their nuclei (Fig. 1a; Fig. S1a,b).

**Figure 1.**
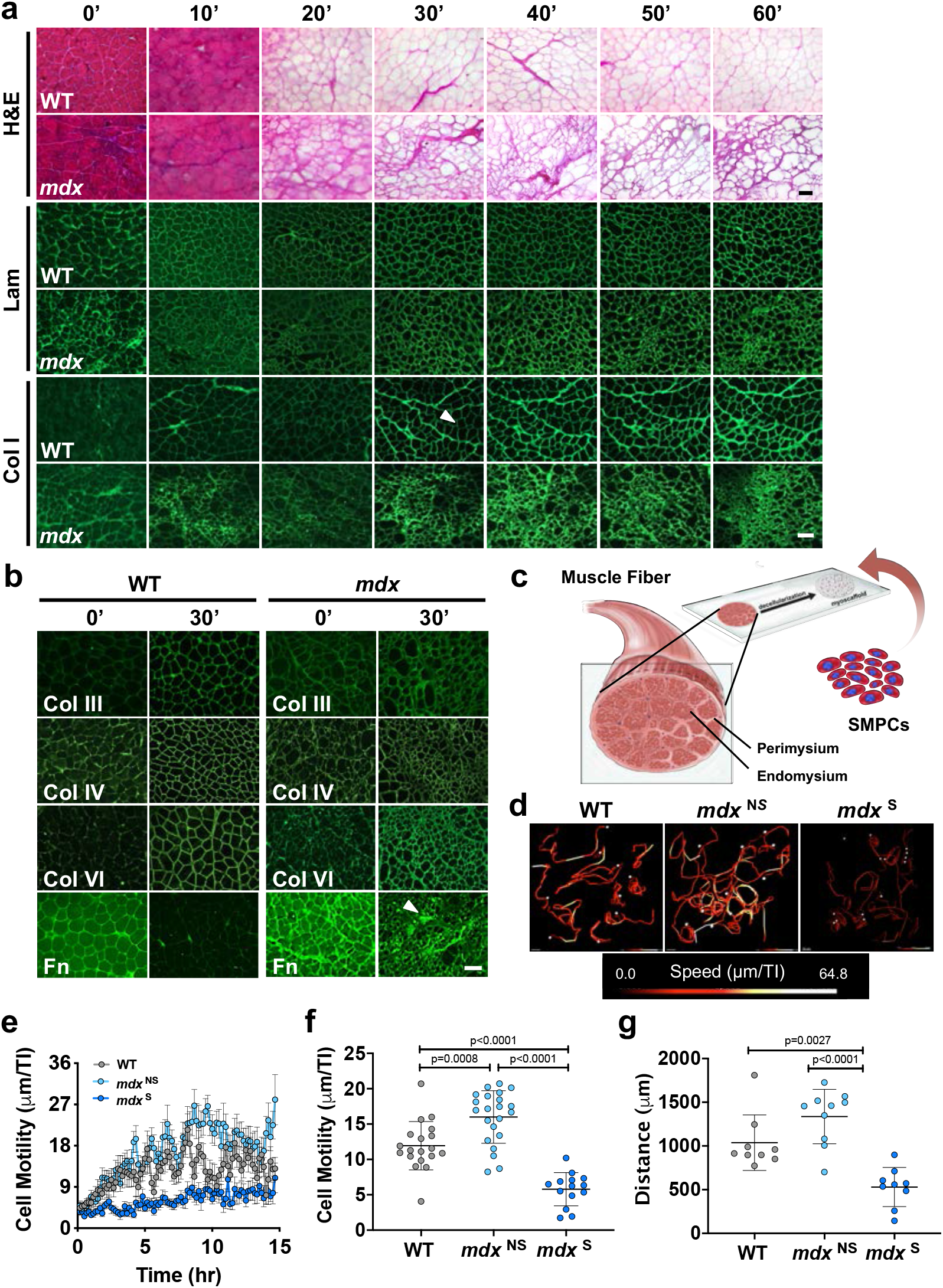
On-slide decellularization of skeletal muscle yields acellular myoscaffolds that recapitulate ECM architecture and composition and support SMPC growth. **a.** Representative images of hematoxylin and eosin (H&E) staining and indirect immunofluorescence (laminin (Lam) and collagen I (Col I)) performed on transverse cryosections from wild-type (WT) and *mdx* quadriceps muscles that were decellularized with 1% SDS solution for the indicated times (10’ to 60’). Non-decellularized (whole) muscle sections were used as controls (0’). The arrowhead on the 30’ Col I WT image indicates the endomysium (n=3-5 independent experiments). Scale bar, 100 μm. **b.** Representative images of intact (0’) and decellularized (30’) muscle sections from WT and *mdx* mice stained using antibodies to collagen types III (Col III), IV (Col IV), and VI (Col VI), as well as fibronectin (Fn). The arrowhead on the 30’ Fn *mdx* image indicates a region of fibrotic scarring (n=3 independent experiments). Scale bar, 100 μm. **c.** Schematic showing the addition of RFP+ SMPCs (WTC-11) onto myoscaffolds derived from decellularized skeletal muscle tissue sections. **d.** Live imaging of RFP+ SMPCs (WTC-11) was used to track cell migration over a 15-hour time period. Cell tracking and heatmap signatures of cell speed show increased SMPC migration rate on *mdx* ^NS^ myoscaffolds, while reduced speeds are observed on *mdx* ^S^ myoscaffolds (n=10-14 cells/tissue; based on observations from n=3 independent experiments). **e.** SMPC motility speeds were calculated based on displacement between individual time points (speed=μm/TI (time interval (TI)=10 minutes)). Graph shows the mean ± SEM. **f.** The average speed (mean ± SD) of SMPCs on the WT, *mdx* ^NS^, and *mdx* ^S^ myoscaffolds were compared by one-way ANOVA (panels e-f: n=14-22 cells/tissue; based on observations from n=3 independent experiments). **g.** Total cell displacement (μm) over the first 15 hours of imaging. Only cells with complete tracks over all frames were included in the analysis (n=9-10 cells/tissue; based on observations from n=3 independent experiments). *P* values reflect analysis by one-way ANOVA, graph shows mean ± SD.

To investigate the effect of decellularization on the localization and abundance of ECM proteins, we performed standard indirect immunofluorescence analysis using antibodies to components of the basement membrane (laminin, collagen IV, and collagen VI) and the interstitial matrix (collagen I, collagen III, and fibronectin). With the exception of fibronectin, decellularization increased exposure of antibody epitopes, leading to improved antibody accessibility and enhanced ECM visualization, enabling improved interrogation of multiple proteins in their native microenvironments (Fig. 1a,b). Of the ECM proteins that were investigated, collagen IV was the only component with decreased expression in *mdx* samples (Fig. 1b). Collagen I was concentrated in the WT and *mdx* perimysium, which is the thickened ECM surrounding bundles of muscle fibers (Fig. 1a). However, collagen I deposition was also evident in the interstitial matrix of the *mdx* endomysium, the layer of connective tissue surrounding individual muscle fibers (Fig. 1a). Fibronectin, which is present in both WT and *mdx* muscle, was largely extracted from WT myoscaffolds after 30 minutes of decellularization, while it was retained in areas of fibrotic scarring in *mdx* samples (Fig. 1b). Increased laminin deposition was also evident in the *mdx* fibrotic scars (Fig. 1a), supporting that the scars likely consist of protein aggregates comprised of collagen and laminin.

### Myoscaffolds retain biological activity that supports cell adhesion and motility

To determine whether myoscaffolds generated using on-slide decellularization retain biological activity, we developed several *in vitro* assays to investigate cell behavior. We previously demonstrated that human pluripotent stem cells (hPSC) can be differentiated to skeletal muscle progenitor cells (SMPCs) that, when enriched for HNK1^-^ ERBB3^+^ NGFR^+^ cell surface receptors, engraft in *mdx* muscle *in vivo* ^34^. However, these SMPCs are still largely inefficient at fusing and regenerating new myofibers *in vivo*. As described above, *mdx* myoscaffolds retain fibrotic scarring that may affect stem cell function or SMPC *in vivo* engraftment. We selected SMPCs to specifically probe the effects of *mdx* ECM on progenitor cells because of their potential for future clinical translation, and to test whether the *mdx* microenvironment negatively influences SMPC function. To investigate SMPC-ECM interaction in real time, we immunolabeled myoscaffolds with a pan-laminin antibody and used hPSCs (WTC-11 line ^35^) expressing a constitutively active red fluorescent protein that localizes to the plasma membrane (mTagRFP inserted in the AAVS1 safe harbor locus to prevent silencing) ^36^. RFP+ hiPSC SMPCs were seeded onto myoscaffolds (Fig. 1c) and images were captured every 10 minutes over a 4-day period using a spinning disc confocal microscope, which enabled high resolution tracking of cell motility and ECM remodeling (Video S1-S4). Videos were analyzed using the Imaris v9.3 software to determine motility behavior and data was collected from all SMPCs in the field of view. Given the heterogeneity observed in the *mdx* ECM due to progression of muscle pathology from early-stage myofiber hypertrophy with limited fibrosis, or scar formation, (*mdx* ^NS^) to late-stage fibrosis with dense scar formation (*mdx* ^S^), we analyzed the behavior of cells in each of these regions separately.

RFP-labeled SMPCs began to settle on the ECM following 10 minutes in culture. SMPCs preferentially adhered to and migrated along the laminin sublayer of myoscaffolds, which represents the basement membrane. On both WT and *mdx* myoscaffolds, cell migration was minimal during the first 2 hours of seeding (average speed=μm/TI (TI: time interval (10 min)); WT: 6.21 μm/TI, *mdx* ^NS^: 6.40 μm/TI, *mdx* ^S^: 6.14 μm/TI). After the first 2 hours, SMPCs cultured on *mdx* ^NS^ myoscaffolds exhibited increased cell motility (Fig. 1d-f; Fig. S1c) and trended toward increased overall migration (distance traveled) (Fig. 1g; Fig. S1d) compared to WT controls. Conversely, SMPCs on *mdx* ^S^ myoscaffolds exhibited a significant reduction in cell motility and migration relative to cells on both WT and *mdx* ^NS^ myoscaffolds (Fig. 1d-g; Fig. S1c-e).

Not only was cell motility different, but cell behavior was also affected between SMPCs cultured on WT and *mdx* myoscaffolds. SMPCs attached to myoscaffolds extended cellular projections that appeared to contract, resulting in mechanical deformation of the myoscaffold at the site of contact and movement of the cell toward the point of attachment (Video S1-S3). Interestingly, we found SMPCs cultured on *mdx* ^NS^ myoscaffolds preferentially circled the circumference of the inner laminin sublayer at a 2.3-fold greater frequency than when on WT myoscaffolds (Fig. S1f,g; Video S2,S4). SMPCs on *mdx* ^S^ myoscaffolds clearly avoided fibrotic scars (Video S3). However, when SMPCs settled into *mdx* ^S^ fibrotic scars, they were unable to deform the ECM, ceased to migrate, and appeared rounded with characteristic apoptotic morphology (Video S3).

We observed that SMPC migration exerted mechanical forces on myoscaffolds that resulted in stretching and deformation of the basement membrane, which preceded remodeling of the laminin sublayer (Video S1, arrows). SMPCs on WT myoscaffolds appeared to degrade laminin at the site of attachment, resulting in a 17% reduction of laminin fluorescence within 30 hours of cell seeding (Fig. S1h-j). While SMPCs on *mdx* myoscaffolds deformed and migrated on the laminin sublayer, migration toward the point of attachment was not always accompanied by ECM remodeling (Video S2). This was especially prominent in fibrotic scars that were not remodeled, as evident by only a 1% reduction in laminin fluorescence after 30 hours in culture, (Fig. S1j). Myoscaffolds enabled the first evaluation of SMPC dynamics in a highly reproducible *ex vivo* system and demonstrated striking differences in the ability of SMPCs to remodel diseased ECM microenvironments.

### Highly crosslinked, stiff *mdx* myoscaffolds are resistant to SMPC-mediated remodeling

Collagen is the most abundant protein in the ECM and provides the main source of extracellular support and force transmission ^37^. In fibrosis, collagen I is significantly upregulated and hypothesized to impair normal cell function ^37^. Given the diminished ECM remodeling and reduced deformation of fibrotic scars within the *mdx* myoscaffolds, we next investigated whether collagen I abundance impairs cell-mediated ECM remodeling and integration. SPMCs were cultured on myoscaffolds for 5 days, followed by detection of collagen I levels using indirect immunofluorescence with anti-collagen I antibodies. To quantify collagen I remodeling, we formulated a remodeling index (RI), calculated as the peak fluorescent intensity of collagen I in the endomysium of myoscaffolds following SMPC culture (5 days) relative to myoscaffolds without SMPCs (media only). RI values close to 1 indicate minimal ECM remodeling, while higher RI values reflect extensive remodeling. SMPCs cultured on WT myoscaffolds degraded collagen I, as indicated by reduced fluorescence (RI, 5.4 ± 2.2) (Fig. 2a,b). SMPCs also integrated throughout the depth of WT myoscaffolds and, through ECM remodeling, extended beyond the former boundaries of the laminin sublayer, thus enabling extensive cell-cell contacts (Fig. 2c). In contrast, collagen I remodeling was inhibited on the *mdx* myoscaffolds (RI, 3.3 ± 1.2) (Fig. 2a-c), which was most pronounced in the fibrotic scars (Fig. S1i). SMPCs appeared rounded and were restricted to the superior surface of the myoscaffolds and limited to the inner laminin sublayer of the basement membrane (Fig. 2a-c).

**Figure 2.**
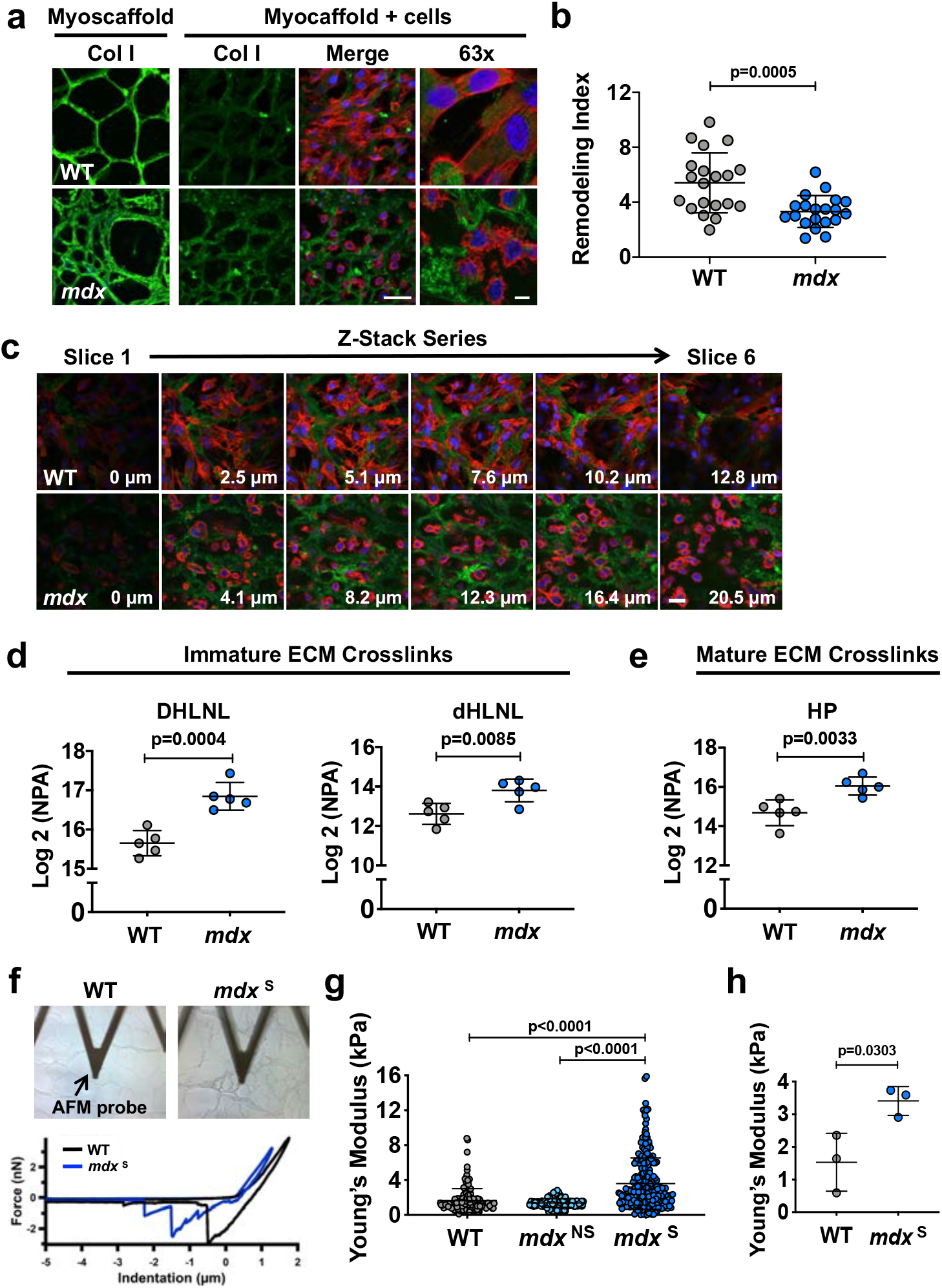
Highly crosslinked, stiff myoscaffolds are resistant to SMPC-mediated remodeling. **a.** Indirect immunofluorescence confocal microscopy of SMPCs (CDMD 1002 cells) cultured for 5 days on WT and *mdx* myoscaffolds (myoscaffold + cells). A decellularized section from each group was not seeded with SMPCs and served as a control (myoscaffold). Sections were stained with antibodies recognizing collagen I (Col I) (green), along with phalloidin as an actin cytoskeleton marker (red) and DAPI as a nuclear marker (blue) (observations from n=5 independent experiments). Scale bars, 25μm (20x), and 8μm (63x). **b.** Graph showing values for the remodeling index (RI: pre/post-remodeling pixel intensity) from WT and *mdx* myoscaffolds stained for Col I (CDMD 1002 cells, n=20 endomysial locations/20x image, results are mean ± SD; based on observations from n=3independent experiments). *P* values reflect analysis by two-tailed unpaired t-test. **c.** Individual tiles from stacked confocal images of SMPCs (CDMD 1002 cells) cultured on WT and *mdx* myoscaffolds. Sections were stained for collagen I (green), phalloidin (red), and DAPI (blue). SMPCs cultured on WT myoscaffolds remodeled and integrated into the ECM, visualized as cells extending into the endomysium and through the myoscaffold thickness. SMPCs cultured on *mdx* myoscaffolds did not integrate into the myoscaffold, localizing adjacent to the basement membrane or on top of the ECM (observations from n=5 independent experiments). Scale bar, 25 μm. **d-e.** Quantification of immature dihydroxy lysinonorleucine (DHLNL) and dehydrohydroxy-lysinonorleucine (dHLNL) crosslinks (**d**), and mature hydroxylyslpyridinoline (HL) crosslinks (**e**) in WT (n=5) and *mdx* (n=5) quadriceps samples. Values are expressed as the Log2 normalized peak area (NPA), mean ± SD. *P* values reflect analysis by two-tailed unpaired t-test. **f.** Images of WT and *mdx* myoscaffolds under the AFM probe during testing and sample traces from the force-separation curves collected from one location on each sample. **g.** Graph showing the Young’s modulus values acquired from a 20 x 20 μm area of the endomysium of a WT myoscaffold (1 biologic sample, n=154 points), along with a *mdx* region without evidence of scars (*mdx* ^NS^)(1 biologic sample, n=256 points) and a *mdx* region with fibrotic scars (*mdx* ^S^)(1 biologic sample, n=240 points) in *mdx* myoscaffolds. Graph showing mean ± SD (n=3 independent experiments). *P* values reflect analysis by one-way ANOVA. **h.** Graph showing the average Young’s modulus values (mean ± SD) collected from 20 x 20 μm areas from WT (n=3 biologic samples, average of 67-154 points/sample) and *mdx* ^S^ (n=3 biologic samples, average of 111-256 points/sample) myoscaffolds. *P* values reflect analysis by two-tailed unpaired t-test.

Collagen fibrils in the ECM are stabilized by the formation of divalent and trivalent intermolecular crosslinks that transmit contractile forces within skeletal muscle ^38^. While crosslinking enzymes are present in healthy tissues, increased crosslinking activity has been implicated in fibrotic diseases and hinders ECM degradation during remodeling ^39^. To determine if collagen crosslinking contributes to the differences in remodeling between WT and *mdx* samples, we used liquid chromatography tandem mass spectrometry (LC-MS/MS) to quantify the abundance of: 1) immature divalent crosslinks (dihydroxy lysinonorleucine (DHLNL), dehydrohydroxylysinonorleucine (dHLNL)) and 2) mature trivalent crosslinks (hydroxylyslpyrodinoline (HP)). A significant increase in both immature and mature crosslinking was observed in the *mdx* muscle relative to control samples (Fig. 2d,e), supporting that collagen crosslinking might contribute to the remodeling barrier in *mdx* myoscaffolds.

Recent mechanical studies have revealed that divalent collagen crosslinking increases tissue compliance, while mature trivalent crosslinking contributes to tissue stiffness ^40^. Immature crosslinks are predominant in areas undergoing active regeneration, while mature crosslinks are associated with established fibrosis ^40-42^. In the *mdx* mouse, skeletal muscle is continuously undergoing degeneration and regeneration, with dense fibrotic scars emerging as a product of failed regeneration. As captured in the live cell time-lapsed videos, SMPCs were unable to deform fibrotic scars within the *mdx* myoscaffolds (Video S3). Based on these observations, coupled with the crosslinking data, we speculate that the immature DHLNL collagen crosslinking in *mdx* scaffolds is predominant in areas of myofiber regeneration, as reflected by increased ECM compliance, while the mature HP collagen crosslinking is enriched in areas of fibrosis, as indicated by increased ECM stiffness. To investigate the ECM mechanical properties, we probed the endomysium of myoscaffolds using atomic force microscopy (AFM) (Fig. 2f). The Young’s modulus, a metric of ECM stiffness, was calculated from the slope of the force-indentation curve obtained during force application. Regions without fibrotic scars were more compliant, with modulus values similar to WT myoscaffolds, while fibrotic scars (*mdx* ^S^) were significantly stiffer (Fig. 2g; Fig. S2a,b). Modulus values for *mdx* ^NS^ regions were also more heterogeneous, consistent with the increase in both mature and immature collagen crosslinking (Fig. 2g). Evaluation of additional fibrotic regions of *mdx* ^S^ myoscaffolds revealed significantly higher average Young’s modulus values relative to controls (Fig. 2h).

### *mdx* myoscaffolds are inherently resistant to laminin remodeling and do not support cell adhesion

During live cell imaging, we observed SMPC-mediated remodeling of laminin in the basement membrane of WT myoscaffolds and in regional-specific areas of *mdx* myoscaffolds within 30 hours of cell seeding (Video S1). Degradation of laminin is a necessary initiating step in myogenesis ^18^. To further evaluate laminin remodeling, SMPCs were again cultured on WT and *mdx* myoscaffolds for 5 days by which time SMPCs had reached confluence. Laminin levels were detected by indirect immunofluorescence using anti-laminin antibodies and relative peak fluorescence intensity was determined, along with the RI for laminin (Fig. 3a,b). Laminin remodeling in the fibrotic *mdx* ECM was inhibited, as revealed by persistent laminin in the basement membrane and a RI close to 1 (RI, 1.25 ± 0.23), relative to the RI of WT controls (RI, 2.35 ± 0.56) (Fig. 3a,b). Interestingly, laminin was far more resistant to remodeling in *mdx* myoscaffolds compared to collagen I, with RI values that were 2.7 fold higher relative to collagen I (Fig. 2b; Fig. 3b).

**Figure 3.**
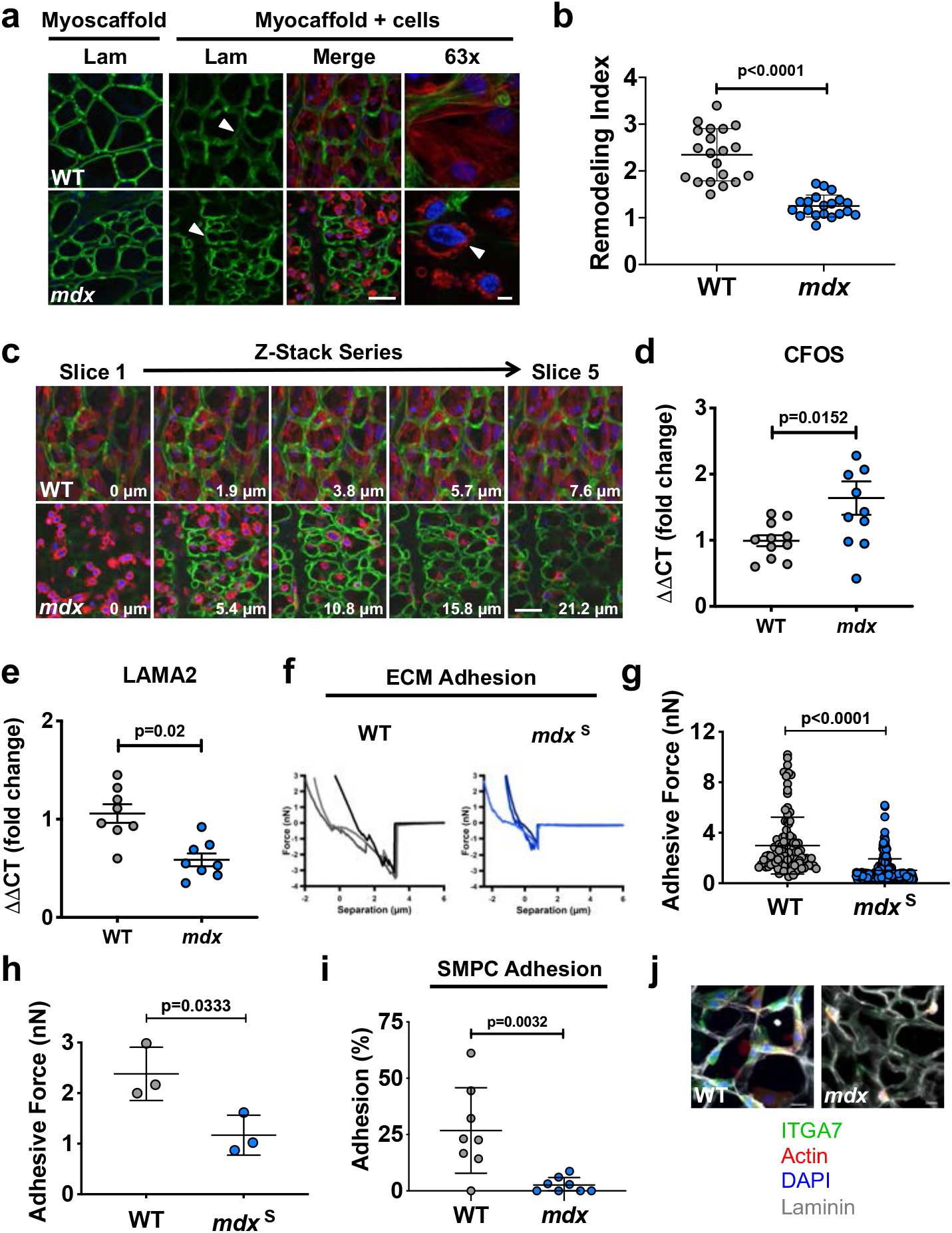
Laminin deposition in *mdx* myoscaffolds impairs SMPC remodeling and adhesion. **a.** Indirect immunofluorescence confocal microscopy of SMPCs (CDMD 1002 cells) cultured for 5 days on WT and *mdx* myoscaffolds (myoscaffold + cells) stained with antibodies recognizing laminin (Lam) (green), along with phalloidin (red) and DAPI (blue). The arrowhead on the WT sample indicates a region where laminin is remodeled, visualized as reduced fluorescence, while the arrowhead on the *mdx* sample indicates a region that is resistant to remodeling, as laminin remains localized in the basement membrane. The arrowhead on the 63x image shows cell membrane blebbing of SMPCs on *mdx* myoscaffolds (observations from n=5 independent experiments). Scale bars, 25 μm (20x), and 8 μm (63x). **b.** Graph showing values for the remodeling index (RI: pre/post-remodeling pixel intensity) from WT and *mdx* myoscaffolds stained for laminin (CDMD 1002 cells, n=20 endomysial locations/20x image, results are mean ± SD; based on observations from n=3 independent experiments). *P* values reflect analysis by two-tailed unpaired t-test. **c.** Individual tiles from z-stacked confocal imaging of SMPCs (CDMD 1002 cells) cultured on WT and *mdx* myoscaffolds stained with laminin (green), phalloidin (red), and DAPI (blue). The first image was taken on the superior portion of the sample, and subsequent images show movement through the thickness of the sample. Scale bar, 25 μm. (observations from n=5 independent experiments) **d-e.** qPCR data showing c-FOS (CFOS) and laminin α2 (LAMA2) expression in SMPCs (CDMD 1002 cells) cultured on WT and *mdx* myoscaffolds for 5 days in proliferation media. Each data point (n=8-11) represents an independent cell culture well from 3 independent experiments (two-tailed unpaired t-test, *p*<0.05). **f.** Representative graphs of the force-separation curves collected during retraction of the AFM probe from a WT and an *mdx* ^S^ myoscaffold. **g.** Graphs showing adhesive force values (mean ± SD) from a 20 x 20 μm area of the endomysium of a WT (1 biologic sample, n=100 points) and *mdx* ^S^ (1 biologic sample, n=256 points) myoscaffold (n=1). *P* values reflect analysis by two-tailed unpaired t-test (n=3 independent experiments). **h.** Graph showing significantly less adhesive force in the *mdx* ^S^ endomysium (n=3 biologic samples, average of 112-256 points/sample), relative to WT samples (n=3 biologic samples, average of 67-154 points/sample). *P* values reflect analysis by t-test. **i.** Graph showing the percent of SMPCs (H9 cells) adhering to WT and *mdx* myoscaffolds (mean ± SD) following exposure to dissociation buffer for 15 minutes, relative to the number of adherent cells before dissociation (cells counted from n=8 3×3 tiled images/group (20x), based on observations from n=4 independent experiments). *P* values reflect analysis by two-tailed unpaired t-test. **j.** SMPCs (CDMD 1002 cells) demonstrated reduced expression of integrin α7 (ITGA7) after 4 hours in culture on *mdx* myoscaffolds, relative to WT controls (observations from n=3 independent experiments). Scale bars, 25 μm.

Z-stacked confocal images were taken of SMPCs cultured on myoscaffolds for 5 days in proliferation media. Consistent with the reduced remodeling activity, we found that SMPCs on *mdx* myoscaffolds were localized primarily on the surface of the tissue with few cells detected deeper within the myoscaffold (Fig. 3c). In contrast, SMPCs were present throughout the WT myoscaffolds, revealed by their integration into the endomysium and throughout the entire Z-plane of the tissue (Fig. 3c). SMPCs on fibrotic regions of *mdx* myoscaffolds were smaller, rounded, and characterized by condensed, perinuclear cortical actin rings and plasma membrane blebbing (Fig. 3a,c). The morphological presentation of SMPCs on *mdx* myoscaffolds is likely reflective of cell stress, which is associated with reduced substrate adhesion ^43-45^, suggesting that the SPMCs are unable to adhere to the *mdx* myoscaffolds. Interestingly, we found increased expression of c-FOS, a gene upregulated in stress-induced cell death ^46,47^, and reduction of laminin α2 expression, the primary ligand for cell attachment to the basement membrane of adult skeletal muscle, in SMPCs cultured on *mdx* myoscaffolds (Fig. 3d,e).

To investigate whether the *mdx* ECM can support cell adhesion, we evaluated the adhesion capacity of *mdx* myoscaffolds by determining the peak rupture force during the retraction phase of AFM. The fibrotic *mdx* myoscaffolds exhibited lower peak breaking forces relative to controls (Fig. 3f-h; Fig. S2a,b). To determine if the decreased adhesive capacity of the *mdx* myoscaffolds affected cell binding, we performed traditional cell adhesion assays in which SMPCs were cultured on WT and *mdx* myoscaffolds for 4 hours, the time required for SMPC adhesion. Only 2% of SMPCs remained attached on the *mdx* myoscaffolds after gentle dissociation, whereas 25% of cells remained adherent to the WT myoscaffolds (*p*<0.05, Fig. 3i), revealing that inherent properties of the *mdx* myoscaffold do not support cell adhesion. Integrin α7 (ITGA7) is the primary integrin responsible for cell adhesion to laminin in adult skeletal muscle. We found that ITGA7 receptors are robustly expressed by SMPCs cultured on WT, but not *mdx*, myoscaffolds (Fig. 3j), leading to the conclusion that a reduction in integrin binding likely contributes to the diminished SMPC adhesion observed on *mdx* myoscaffolds. In addition, we observed downregulation of genes associated with cell adhesion and migration (*L1CAM, CLSTN2, ITGA4, ITGA6, CDH15, and NCAM*), and skeletal muscle development and maturation (*MYH7B, DMD, DCX, TNC, MIR133B*) in SMPCs cultured on *mdx* myoscaffolds for 5 days, demonstrating that impaired adhesion to the ECM has detrimental effects on cell function (Fig. S2c; Table S1).

### Overexpression of laminin-binding complexes induces a compensatory matrisome

In skeletal muscle, the composition and organization of the ECM is influenced by expression of ECM receptors at the sarcolemma ^19,48-50^. We interrogated how loss of the dystrophin-glycoprotein complex, one of the primary cell surface receptors in skeletal muscle that binds to laminin in the basement membrane, affects *mdx* ECM composition by determining the matrisome, which consists of core ECM proteins, including glycoproteins, collagens, and proteoglycans, in addition to ECM affiliated proteins ^51^. Although prior studies have reported the ECM composition for WT and *mdx* muscles ^52-54^, traditional mass spectrometry protocols fail to capture the majority of fibrillar ECM proteins and thus a comprehensive analysis of the skeletal muscle matrisome is challenging. We used a previously established method for insoluble ECM characterization to achieve improved analysis of mouse skeletal muscle ^55^. To determine if overexpression of cell adhesion complexes in the muscle cell membrane would influence ECM composition, we also investigated *mdx* muscle genetically engineered to overexpress sarcospan. Sarcospan is a well-described transmembrane protein that acts as a scaffold to stabilize laminin-binding transmembrane complexes at the muscle cell surface ^56,57^. Overexpression of sarcospan in *mdx* mice (*mdx* ^TG^) ameliorates muscular dystrophy by increasing expression of integrins at the sarcolemma, which restores muscle fiber attachment to laminin in the basement membrane and protects the muscle from contraction-induced injury ^24,25,58-62^. We interrogated the effect of sarcospan overexpression in *mdx* muscle on ECM composition by determining its matrisome and analyzing it relative to controls ^51^.

We identified a total of 1,679 proteins from all three genotypes, including 57 matrisome and matrisome-associated proteins (Table S2). Principle component analysis revealed distinct clustering of the three sample types, revealing a divergence of the *mdx* ^TG^ matrisome from both WT and *mdx* (Fig. 4a). In general, ECM proteins were less abundant in WT compared to *mdx* samples (Fig. S3a,b; Table S2). Interestingly, integrin α7 (ITGA7) expression was significantly upregulated in *mdx* ^TG^ relative to *mdx* and WT muscle (Fig. 4b). Our group has previously shown that increased integrin α7 (Itga7) at the sarcolemma of *mdx* ^TG^ muscle is required to ameliorate pathology and restore muscle attachment to laminin, which supports the proteomic data ^24,25,58-62^.

**Figure 4.**
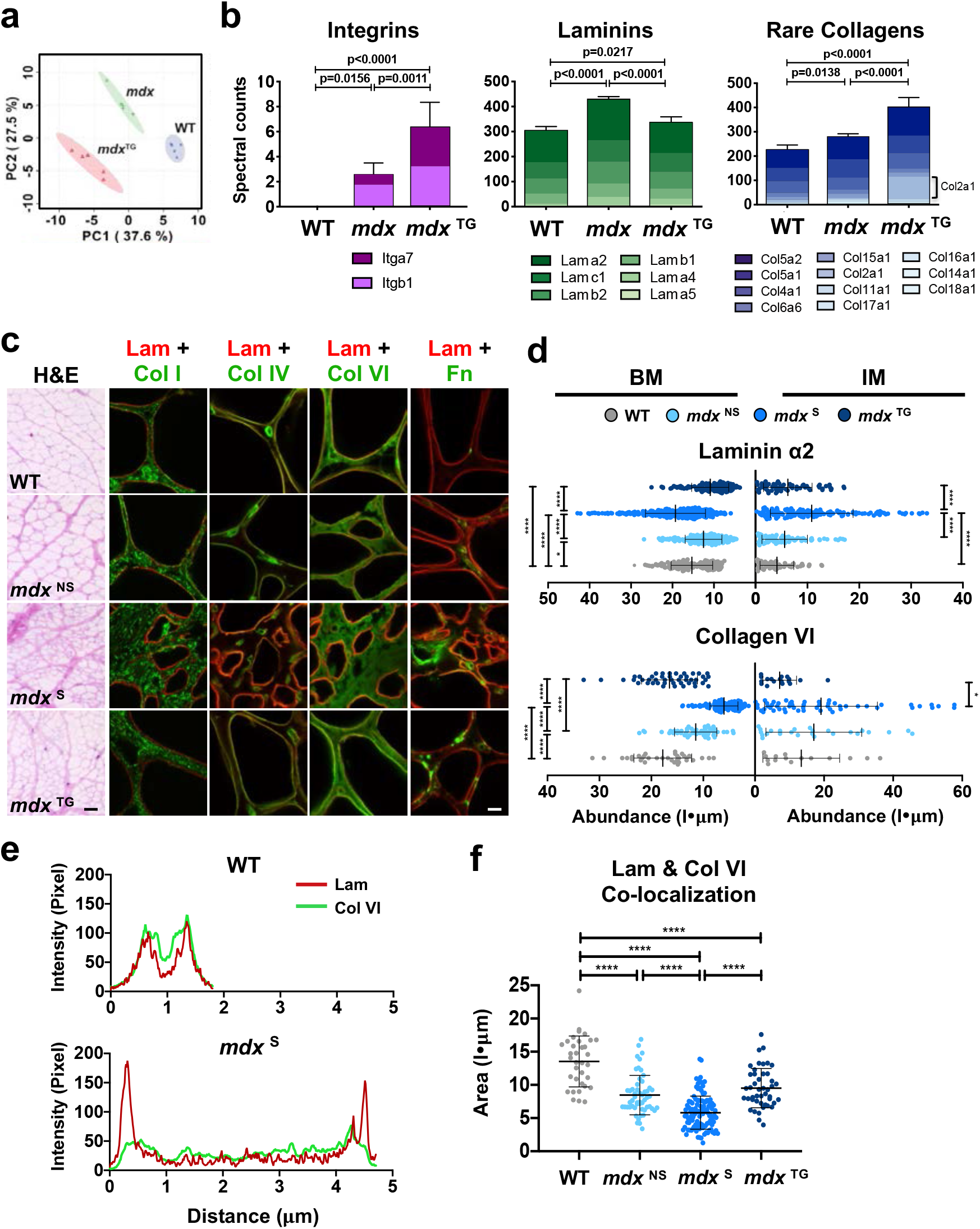
Overexpression of laminin-binding complexes modulates the matrisome and alters ECM organization towards a compensatory physiological phenotype. **a.** Principle component analysis from ECM focused proteomics reveals distinct clustering of each phenotype, with the *mdx*:SSPN-Tg (*mdx* ^TG^) diverging from both the WT and *mdx* samples (n=5 samples/group). **b.** Column graphs showing the abundance of integrins, laminins, and rare collagens not normally expressed in adult skeletal muscle from WT, *mdx*, and *mdx* ^TG^ samples (mean ± SD). *P* values reflect analysis by one-way ANOVA. **c.** Representative images from myoscaffolds stained with H&E, along with immunofluorescent analysis (IFA) of myoscaffolds co-stained for laminin α2 (Lam) (red) with collagen I (Col I), IV (Col IV), VI (Col VI), and fibronectin (Fn) (green), respectively, from WT, *mdx* ^NS^, *mdx* ^S^, and *mdx* ^TG^ samples (selected images from n=4 independent experiments). Scale bars, 100μm (H&E) and 8μm (IFA). **d.** Graphs showing the abundance (pixel intensity x um (I•μm)) of laminin α2 and collagen VI in the basement membrane (BM) and interstitial matrix (IM) of WT, *mdx* ^NS^, *mdx* ^S^, and *mdx* ^TG^ ECM (mean ± SD). Between group differences were analyzed by one-way ANOVA. *P* values are as follows: * = *p<0.05*, ** = *p<0.01*, *** = *p<0.001*, **** = *p<0.0001*. (n=20-40 measurements/group for each component except laminin, where n=80-150 measurements/group) **e.** Representative plot profiles of the pixel intensity of laminin α2 (red) and collagen VI (green), as measured across the width of one endomysial location from confocal images of one WT and *mdx* myoscaffold. The peaks in the red channel represent laminin α2 in the basement membrane. The pixel intensity of laminin α2 in the basement membrane of the *mdx* endomysium is approximately double of that observed in the WT endomysium while the intensity of collagen VI is reduced by half. **f.** Graph showing the relative stoichiometry between laminin α2 and collagen VI in the basement membrane of WT, *mdx* ^NS^, *mdx* ^S^, and *mdx* ^TG^ ECM. Between group differences were analyzed by one-way ANOVA. *P* values are as follows: * = *p<0.05*, ** = *p<0.01*, *** = *p<0.001*, **** = *p<0.0001*.

We found that approximately 60% of matrisome and matrisome-associated proteins were upregulated in *mdx* compared to *mdx* ^TG^ or WT, including several laminins, annexins, periostin, and nidogen (Table 2; Fig. 4a; Fig. S3a,b). Laminin α2 and β1 (Fig. 4b) were upregulated in *mdx* samples, consistent with immunofluorescence analysis of the myoscaffolds (Fig. 1a). However, overexpression of sarcospan reduced expression of all laminin isoforms in the *mdx* ^TG^ samples relative to *mdx* controls (Fig. 4b). In fact, the laminin profile of *mdx* ^TG^ muscle was more similar to WT.

Many collagens expressed in low abundance in WT skeletal muscle, including collagen V and XI, were significantly elevated in *mdx* ^TG^ samples (Fig. 4b; Fig. S3a-c). As collagens function to transmit forces from the muscle to the bone, the increased abundance of collagens in the *mdx* ^TG^ ECM may reflect a compensatory strategy to support improved force transmission. Interestingly, galectin 1 (Lgals1), which promotes cell migration and growth and aids in the conversion of stem cells to myogenic cells during muscle repair and regeneration ^63-66^, was also significantly upregulated in *mdx* ^TG^ samples relative to both WT and *mdx* (Fig. S3b). Overall, the matrisome profile of the *mdx* ^TG^ samples exhibits a compensatory remodeling phenotype.

### Loss of myofiber adhesion in *mdx* muscle causes ECM disorganization that is corrected by overexpression of integrin complexes

Based on previous studies revealing a role for laminin-binding membrane receptors in organizing the ECM ^48^, we sought to investigate the ECM organization of *mdx* and *mdx* ^TG^ muscle. Analysis of H&E staining of whole and decellularized *mdx* ^TG^ muscle sections revealed improved ECM architecture relative to *mdx* muscle, resembling that observed in WT samples (Fig. 4c; Fig. S3d, S4a). Interestingly, despite the improvement in muscle pathology, the *mdx* ^TG^ was also characterized by increased ECM deposition, observed as thickening of the endomysium and perimysium (Fig. 4c). To determine the localization of proteins within the ECM sublayers, myoscaffolds were analyzed after indirect immunofluorescence detection of laminin α2 (red fluorescence), co-stained with antibodies to the following ECM components: fibronectin, collagen I, IV, and VI (green fluorescence) (Fig. 4c; Fig. S3d). These proteins were selected to represent both the basement membrane (laminin α2, collagen IV, and collagen VI) and interstitial matrix sublayers (fibronectin and collagen I). Using line scan data generated with Image J software, we created a MATLAB algorithm to determine the abundance of each protein within the basement membrane and interstitial matrix between two adjacent myofibers (Fig. S4b).

ECM organization in *mdx* myoscaffolds was evaluated in regions with (*mdx* ^S^) and without (*mdx* ^NS^) fibrotic scars. Reduced deposition of laminin, collagen IV and VI characterized active regeneration in the *mdx* ^NS^ basement membrane, suggestive of thinning, while fibronectin was expanded in both the basement membrane and interstitial matrix (Fig. 4d; Fig. S4c). In contrast, laminin was significantly expanded in the *mdx* ^S^ basement membrane while collagen IV and VI were reduced, revealing a thickening of this sublayer (Fig. 4d; Fig. S4c). Furthermore, fibronectin, normally restricted to the interstitial matrix, was mislocalized and highly expressed in the basement membrane in fibrotic scars (Fig. S4c). In addition to increased deposition in the basement membrane, laminin and fibronectin were also present in the interstitial matrix in *mdx* ^S^ fibrotic scars. These data suggest that, as chronic injury progresses from active regeneration to necrosis with fibrosis, laminin deposition accumulates within the tissue creating a thickened basement membrane.

Line scan analysis of *mdx* ^TG^ myoscaffolds revealed ECM organization similar to that observed in regenerating regions of *mdx* myoscaffolds, including decreased expression of collagen IV and laminin in the basement membrane (Fig. 4d; Fig. S4c). Interestingly, collagen VI was restored to WT levels. The most remarkable finding was the absence of laminin thickening in the basement membrane (Fig. 4d), supporting that laminin organization is retained in both regenerating *mdx* and *mdx* ^TG^ ECM.

Collagen VI has been identified as a key component responsible for regulation of mechanical properties of the satellite cell niche ^67^. A decrease in collagen VI has been linked to impaired muscle regeneration and decreased satellite cell self-renewal capacity following injury ^67,68^. Analysis of the stoichiometry of laminin and collagen VI in the basement membrane of WT myoscaffolds reveals almost complete overlap of the two components (Fig. 4e,f). Conversely, expansion of laminin in the basement membrane of fibrotic *mdx* myoscaffolds is accompanied by a decrease in collagen VI, disrupting the normal stoichiometry of the two components necessary for satellite cell self-renewal (Fig. 4e,f). Overexpression of sarcospan improved the laminin to collagen VI stoichiometry (Fig. 4e,f).

### Laminin α2 organization determines SMPC adhesion and remodeling capacity

SMPCs cultured on *mdx* ^TG^ myoscaffolds effectively remodeled laminin, as demonstrated by significantly higher RI values relative to *mdx* controls (Fig. 5a,b; Fig. S5a-d, S6a,b). In contrast, collagen I remodeling was less effective in *mdx* ^TG^ scaffolds (Fig. 5a,b; Fig. S5a, S6c,d), possibly due in part to the 10% and 25% increase in collagen I expression relative to WT and *mdx* samples, respectively (Fig. S3c; Table 2). Consistent with reduced collagen remodeling, we observed an increase in both immature and mature collagen crosslinks in *mdx* ^TG^ muscle, similar to the levels observed in *mdx* samples (Fig. 5c). Interestingly, the abundance of immature crosslinks was elevated even above the levels observed in *mdx* muscle.

**Figure 5.**
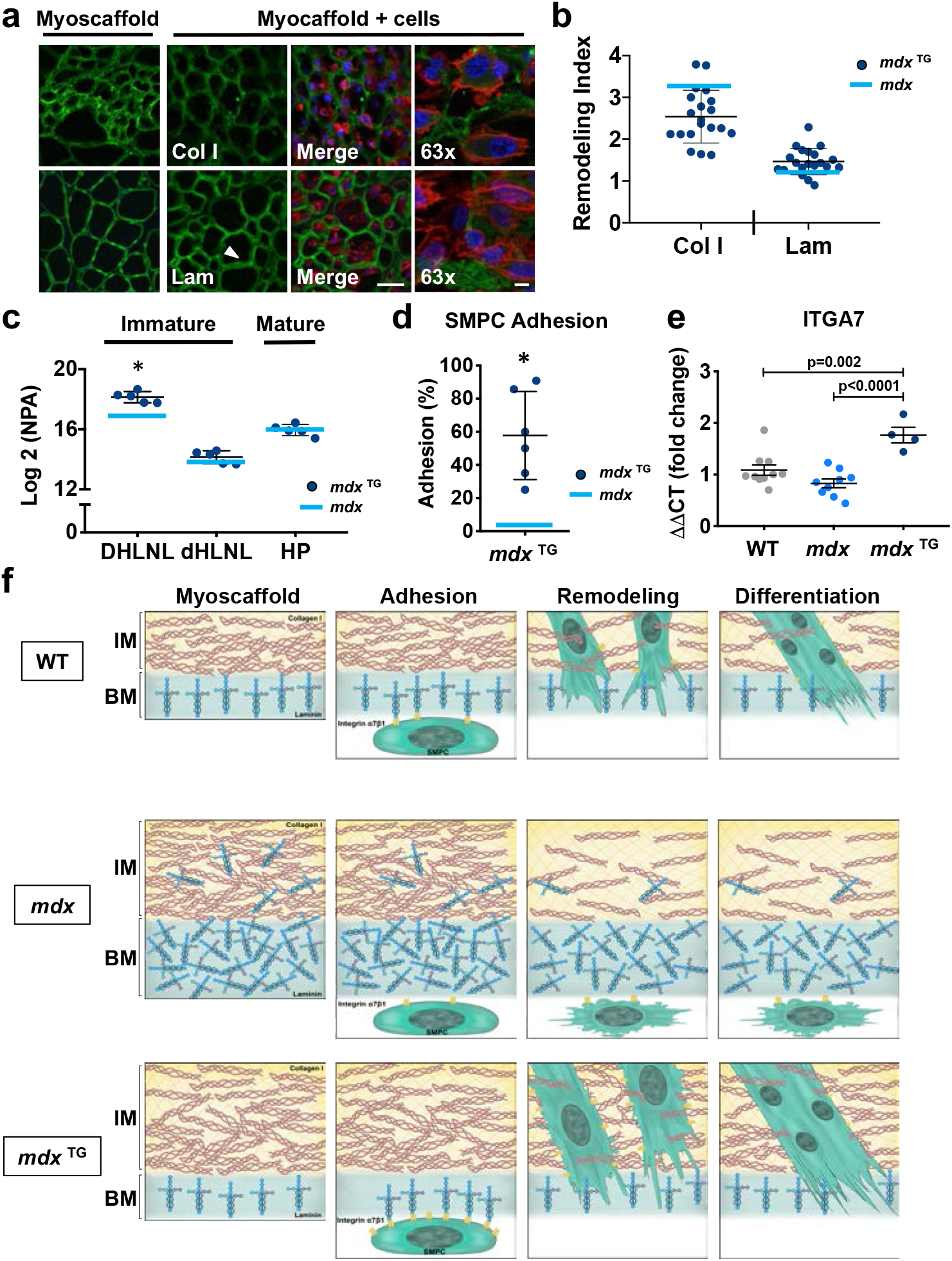
Expansion of laminin in the *mdx* basement membrane is a predictor of impaired SMPC adhesion and ECM remodeling. **a.** Indirect immunofluorescence confocal microscopy of SMPCs (CDMD 1002 cells) cultured for 5 days on *mdx* ^TG^ myoscaffolds (myoscaffold + cells) stained with antibodies against collagen I (Col I) or laminin (Lam) (green), along with phalloidin (red) and DAPI (blue). The arrowhead indicates a region of laminin remodeling, indicated by a reduction in the localization of laminin in the basement membrane (n=5 independent experiments). Scale bars, 25μm (20x), and 8μm (63x). **b.** Calculation of the remodeling index (RI: pre/post-remodeling pixel intensity) from *mdx*^TG^ myoscaffolds stained for Col I and Lam. For reference, the blue bar on the graph represents the mean RI value from SMPCs cultured on *mdx* myoscaffolds (n=20 endomysial locations/20x image, results are mean ± SD; based on observations from n=3 independent experiments). *P* values reflect analysis two-tailed unpaired t-test (* = *p<0.05*). **c.** Quantification of immature dihydroxy lysinonorleucine (DHLNL) and dehydrohydroxy-lysinonorleucine (dHLNL) crosslinks and mature hydroxylyslpyridinoline (HL) crosslinks in *mdx* ^TG^ samples (n=5). Values are expressed as the Log2 normalized peak area (NPA), mean ± SD. The blue bar represents the Log2 (NPA) value from *mdx* samples. *P* values reflect analysis by two-tailed unpaired t-test. **d.** SMPCs (H9 cells) cultured on *mdx* ^TG^ myoscaffolds exhibited improved ECM adhesion following exposure to dissociation buffer. The blue bar represents the mean value from SMPCs cultured on *mdx* myoscaffolds. Cells counted from n=8 3×3 tiled images/group (20x), based on observations from n=5 independent experiments. *P* values reflect analysis by two-tailed unpaired t-test (* = *p<0.05*). **e.** Graph showing the fold change in gene expression of integrin α7 (ITGA7) from SMPCs (CDMD 1002 cells) cultured on WT, *mdx*, and *mdx* ^TG^ myoscaffolds for 5 days (mean ± SD). Each data point (n=4-10) represents an independent cell culture well from n=3 independent experiments. *P* values reflect analysis by one-way ANOVA. **f.** Schematic representation of the abundance and organization of laminin and collagen I in the basement membrane (BM) and interstitial matrix (IM) of WT, *mdx*, and *mdx* ^TG^ myoscaffolds, along with SMPC adhesion, remodeling and differentiation. While SMPCs cultured on WT myoscaffolds adhered to laminin in the basement membrane, permitting successful remodeling of both laminin and collagen I and supporting SMPC differentiation, laminin disorganization in the *mdx* myoscaffolds inhibited SMPC adhesion, blocking laminin remodeling necessary for downstream differentiation. Integrin-priming in the *mdx* ^TG^ muscle restored laminin organization that we hypothesize stimulates the upregulation of integrin α7 expression in SMPCs, permitting robust SMPC adhesion and laminin remodeling required for downstream differentiation.

Despite the diminished capacity to remodel collagen, SMPCs exhibited improved, WT-like cell morphology on *mdx* ^TG^ myoscaffolds, with elongated cells integrated within the myoscaffold (Fig. 5a). Surprisingly, SMPC adhesion to *mdx* ^TG^ myoscaffolds was greater than *mdx* samples (Fig. 5d; Fig. S6e). Similar to WT, we found that SMPCs remained adherent to the inner laminin sublayer of *mdx* ^TG^ myoscaffolds (Fig. S6f). Consistent with increased adhesion, we observed a significant upregulation in *ITGA7* expression in SPMCs cultured on *mdx* ^TG^ myoscaffolds, suggesting that the *mdx* ^TG^ ECM retains properties associated with increased integrin α7 binding that, in turn, regulate SMPC gene expression (Fig. 5e; Fig. S6g). We conclude that elevation of integrin α7β1D in *mdx* ^TG^ myofibers functions to organize laminin in the ECM in a manner that improves integrin-mediated binding of SMPCs in the myoscaffold assay (Fig. 5f). In contrast, *mdx* myofibers lack robust expression of laminin-binding adhesion complexes leading to disorganization of the basement membrane that is characterized by loss of accessible laminin binding sites required for SMPC adhesion and remodeling (Fig. 5f).

### Myotube differentiation requires laminin remodeling and is inhibited by fibrosis

The downregulation of genes associated with cell differentiation and skeletal muscle maturation following SMPC culture on *mdx* myoscaffolds (Fig. S2c) suggest that inherent factors in the ECM influence SMPC gene expression in a manner that may block stem cell differentiation. To investigate the effect of the ECM on cell differentiation, SMPCs were cultured (5 days) in proliferation media on myoscaffolds prepared from WT, *mdx*, and *mdx* ^TG^ muscle, followed by differentiation (5 days). On the WT samples, SMPCs fused into healthy, elongated myofibers that integrated into the myoscaffolds, which was less evident on *mdx* samples (Fig. 6a; Fig. S7a,b). Quantification of SMPC myotube differentiation reveals significant impairment on *mdx* myoscaffolds relative to WT (Fig. 6b). SMPCs extensively remodeled laminin in *mdx* ^TG^ myoscaffolds and exhibited robust fusion efficiency that was identical to WT myoscaffolds (Fig. 6b). Higher magnification images revealed that the differentiated myotubes on *mdx* ^NS^ myoscaffolds are confluent across the myoscaffold and are associated with extensive laminin remodeling (Fig. 6c,d). Notably, myotubes on *mdx* ^S^ myoscaffolds were confined to the inner laminin sublayer of the basement membrane and laminin degradation was restricted to regions of direct cell-ECM contact (Fig. 6c,d). Furthermore, cell fusion was only evident in regions with concomitant laminin degradation; supporting that laminin remodeling is required for differentiation (Fig. 6c,d).

**Figure 6.**
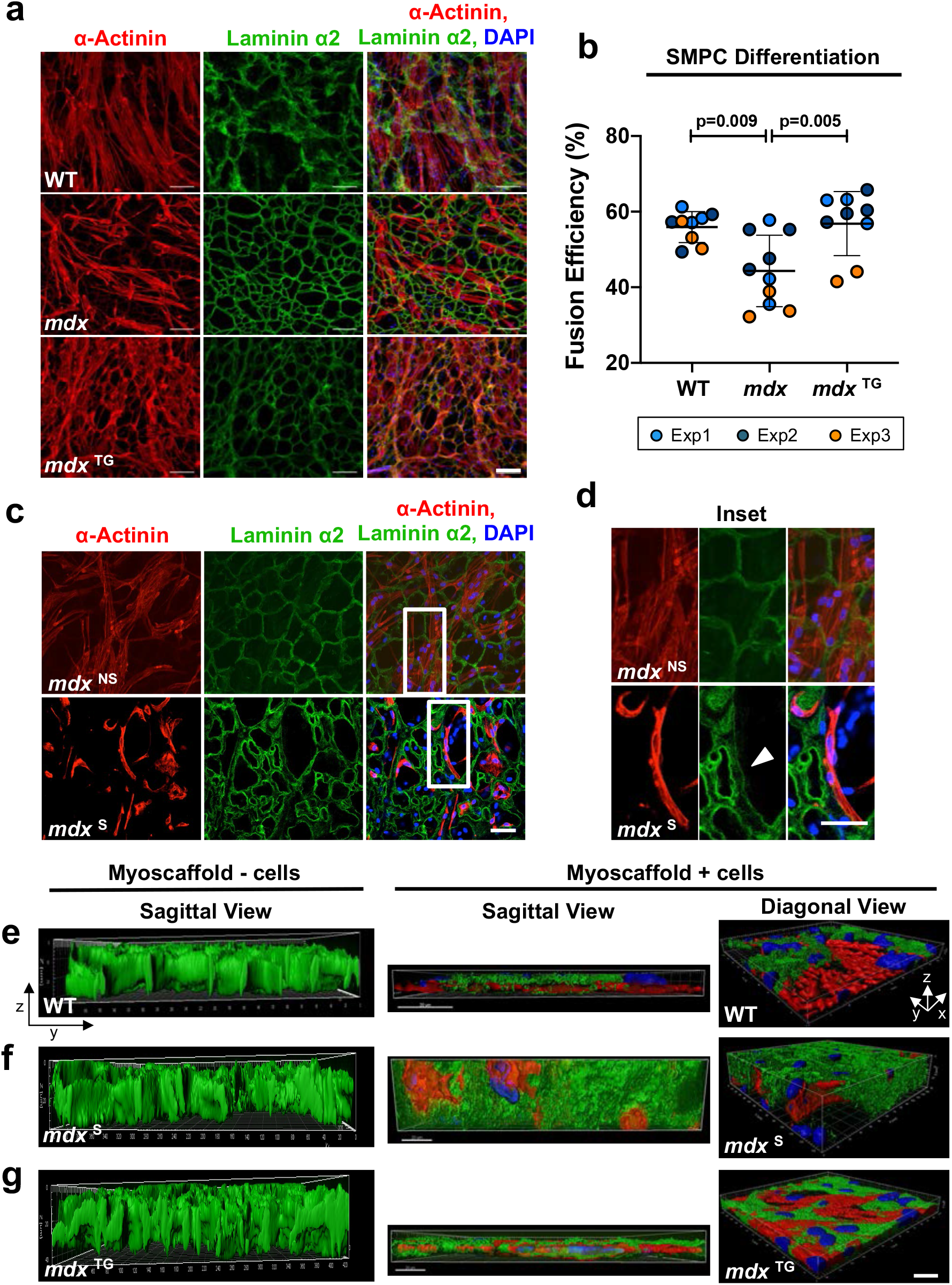
Differentiation requires laminin remodeling and is inhibited by fibrosis. **a-b.** SMPC differentiation on WT, *mdx*, and *mdx* ^TG^ myoscaffolds show representative ability to remodel laminin α2 (green) and fuse to form myotubes. Fusion efficiency was calculated as the percentage of nuclei (DAPI, blue) in myotubes (≥ 3 nuclei per α-Actinin+ cell, red) relative to all nuclei per field of view (mean ± SD). Scale bar, 100 μm. *P* values reflect analysis by one-way ANOVA, *\* = p<0.05* (Fusion efficiency on n=9-10 separate tissues from n=3 independent experiments were quantified, using cell lines CDMD 1002 (Exp1) and H9 (Exp2 & Exp3)). **c-d.** Confocal optical sections show myotubes attach and locally remodel laminin α2 in *mdx* ^LS^ matrix regions, whereas myotubes more readily degrade and integrate with laminin α2 in *mdx* ^ES^ regions. Scale bar, 50 μm. Selected images from H9 cells, based on observations from n=4 independent experiments. **e-g.** Compiled z-stack confocal images exported to Imaris analysis software show myotube integration into WT (**e**), *mdx* (**f**), and *mdx* ^TG^ (**g)** myoscaffolds from diagonal (X-Y-Z) and sagittal (X-Z) views (Laminin α2 (green), α-Actinin (red), and DAPI (blue). Control myoscaffolds that had not been cultured with cells served as controls (myoscaffold – cells) to show the striking reduction in ECM thickness observed in WT and *mdx* ^TG^ samples, relative to *mdx* scaffolds that were resistant to degradation. Scale bar, 20 μm; grid display in diagonal view shows 10 μm increments. Selected images using H9 cells, based on observations from n=4 independent experiments.

Z-stacked confocal images reveal the dramatic effect of SMPC remodeling of WT myoscaffolds, visualized as the substantial reduction in myoscaffold thickness (Fig. 6e). Profound resistance of *mdx* ^S^ ECM to remodeling is evident by the preserved bulk of *mdx* myoscaffolds, even after 10 days of culture with SMPCs (Fig. 6f). The improved capacity for SMPC adhesion, facilitated by the integrin-mediated assembly of the basement membrane, is strikingly evident in the remodeling of *mdx* ^TG^ myoscaffolds (Fig. 6g).

To confirm that the improvement in fusion efficiency observed on *mdx* ^TG^ myoscaffolds is due to inherent properties of the ECM and not due to the reduction in muscle injury and regeneration previously reported in *mdx* ^TG^ mice ^24,25,58-62^, we induced muscle injury in WT, *mdx*, and *mdx* ^TG^ mice and evaluated SMPC adhesion and fusion efficiency on myoscaffolds derived from injured muscles. To induce injury and regeneration, tibialis anterior (TA) muscles were injected with barium chloride (BaCl_2_) ^69^. Muscle regeneration at 7 days was confirmed with the presence of centrally nucleated myofibers (Fig. S8a,b). Myoscaffolds were generated from TA muscles at 7 days post-injury (Fig. S8c) and cultured with SMPCs. All myoscaffolds derived from BaCl_2_ treated muscles supported improved SMPC adhesion compared to myoscaffolds from untreated muscles (Fig. 3i, 5d, 7b, S6e). However, consistent with our previous findings, there was a significant reduction in SMPC adhesion to *mdx +* BaCl_2_ myoscaffolds compared to WT and *mdx* ^TG^ *+* BaCl_2_ myoscaffolds (Fig. S8d,e). Sparse, thin myotube formation and impaired fusion efficiency were observed on *mdx +* BaCl_2_ myoscaffolds, compared to robust myotube formation following differentiation on WT and *mdx* ^TG^ *+* BaCl_2_ myoscaffolds (Fig. S8f,g).

**Figure 7.**
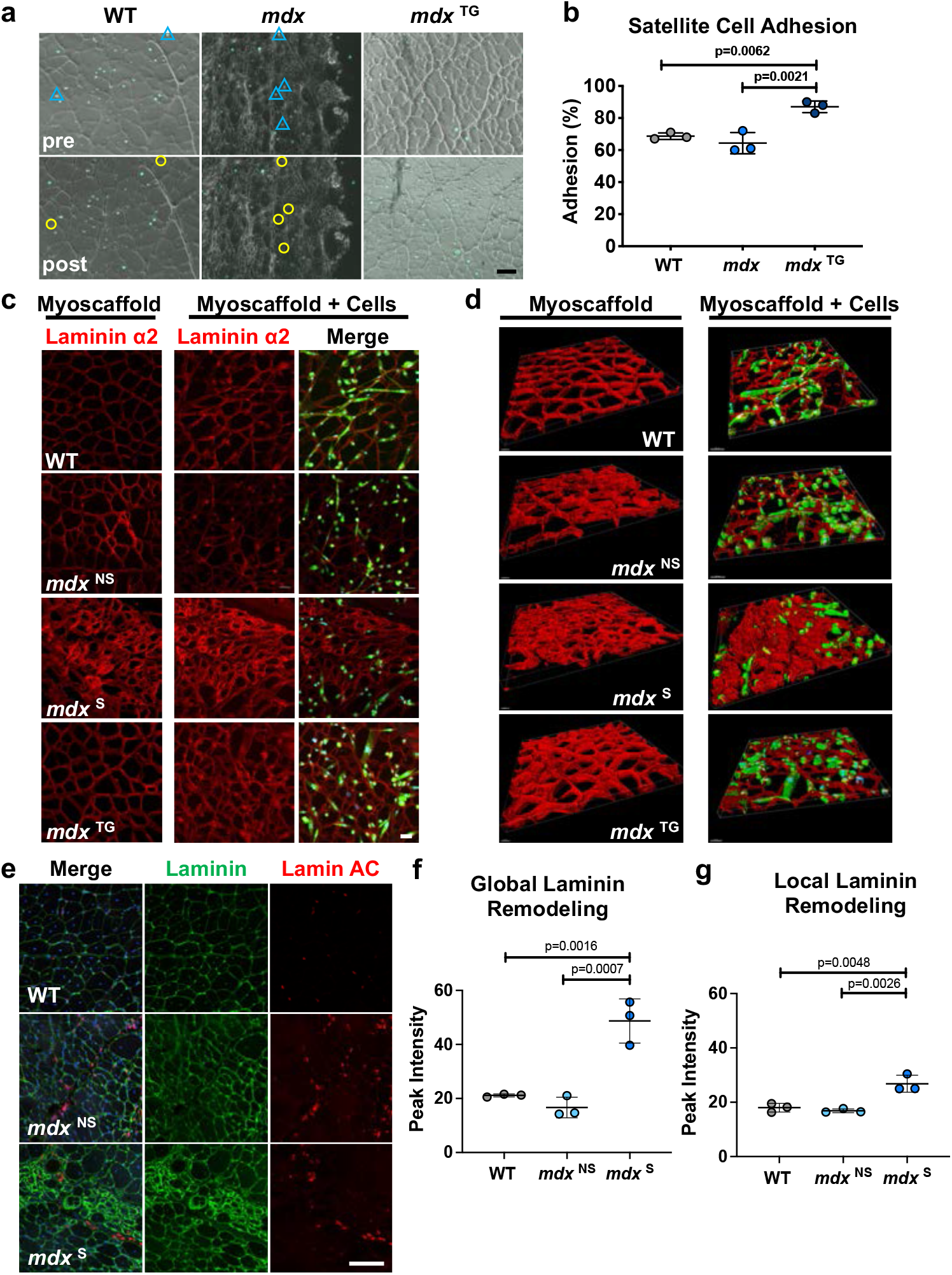
Diminished *in vitro* satellite cell function on fibrotic *mdx* ^F^ myoscaffolds is recapitulated with limited SMPC engraftment *in vivo*. **a.** Phase contrast images showing satellite cells (green fluorescence) on WT, *mdx*, and *mdx* ^TG^ myoscaffolds, both before (pre) and after (post) exposure to dissociation buffer. Blue triangles indicate cell location prior to dissociation and yellow circles indicate the same region where cells were subsequently absent following dissociation. Scale bar, 100 μm. (representative images from n=3 samples/group) **b.** Graph showing the percentage of satellite cells adhering to WT, *mdx*, and *mdx* ^TG^ myoscaffolds following exposure to dissociation buffer (mean ± SD). Cells were counted from images of the entire myoscaffold (stitched images (10x), n=3 samples/group). In general, satellite cells adhered more strongly to myoscaffolds, compared to SMPCs (Fig. 3i, 5d). A similar trend in adhesion is observed with both cell types, with cells on the *mdx* ^TG^ myoscaffolds adhering best, followed by those on WT and then *mdx* myoscaffolds. *P* values reflect analysis by one-way ANOVA. **c.** Confocal images of ZsGreen fluorescent satellite cells cultured on WT, *mdx* ^NS^*, mdx* ^S^, and *mdx* ^TG^ myoscaffolds for 4 days in differentiation media and stained with laminin α2 (red). Myoscaffolds not seeded with cells were maintained in differentiation media and served as controls. Scale bar, 50 μm. Representative images from n=3 samples/group. **d**. Compiled z-stack confocal images exported to Imaris analysis software show limited cell integration into *mdx* ^S^ myoscaffolds from a diagonal (X-Y-Z) view (laminin α2 (red), DAPI (blue)). Scale bar, 30-50 μm. **e.** Representative images from SMPC injected WT C57-NSG and *mdx*-NSG mice, stained for laminin (green), lamin AC (red), and DAPI (blue). SMPCs in WT and regions of the *mdx* muscle without laminin scars (*mdx* ^NS^) integrated throughout the tissue, while cells were unable to penetrate thickened laminin in fibrotic scars (*mdx* ^S^) (n=3 directed differentiations, 2 mice engrafted/differentiation (WT and *mdx*), n=6 mice total). Scale bar, 100 μm. **f-g.** Quantification of laminin intensity from a 10x image of each injected muscle reveals significant laminin deposition in *mdx* ^S^ regions, relative to WT and *mdx* ^NS^ (mean ± SD, n=3/group). *P* values reflect analysis by one-way ANOVA. **(f).** Quantification of laminin intensity in the basement membrane bordering engrafted cells reveals similar laminin intensity as that observed globally in WT and *mdx* ^NS^ muscle, whereas we saw roughly a 45% reduction in laminin intensity surrounding engrafted cells in *mdx* ^S^ regions (**g**) (mean ± SD, n=3/group). *P* values reflect analysis by one-way ANOVA.

### Myoscaffolds reproduce mouse satellite cell behavior and are predictive of engrafted hPSC SMPCs laminin remodeling *in vivo*

To further evaluate the applicability of myoscaffolds, we investigated primary mouse satellite cell behavior on mouse myoscaffolds, and also evaluated *in vivo* engraftment dynamics of hPSC derived SMPCs in wild-type C57bl/10-NOD *scid* gamma (NSG) and dystrophic *mdx*-NSG mice. To investigate satellite cell adhesion, we used a Pax7-Cre Rosa26-ZsGreen mouse model that constitutively fluoresces after tamoxifen induced Cre-recombination, which enabled quantitative live imaging of satellite cells from the same myoscaffold regions before and after cell dissociation. Satellite cells were sorted (Cd45-Ter119-ZsGreen+) and immediately seeded on WT, *mdx*, and *mdx* ^TG^ myoscaffolds. Satellite cells adhered to the myoscaffolds by 16 hours, at which time we used gentle dissociation buffer to test adherence to the different myoscaffolds. Mouse satellite cells were more adherent to the myoscaffolds relative to SMPCs, and relatively few cells dissociated using the dissociation parameters that were effective for SMPCs. Thus, we increased the EDTA concentration, NaCl osmolality, and time in buffer to increase stringency of the assay. Using these conditions, we found 64% of murine satellite cells were retained on *mdx* myoscaffolds, whereas 87% were retained on *mdx* ^TG^ myoscaffolds (Fig. 7a,b).

We also set out to determine whether satellite cells would differentiate on WT, *mdx*, and *mdx* ^TG^ myoscaffolds. Primary satellite cells were expanded as myoblasts prior to seeding on myoscaffolds, and differentiated to myotubes for 4 days. Mouse satellite cells were less proliferative and appeared to be at an earlier stage of matrix remodeling, as indicated by an increase in the width of the endomysium, but no change in laminin fluorescence (Fig. 9a, Fig. 7c,d). However, similar to SMPCs, we found that murine satellite cells avoided fibrotic regions of *mdx* myoscaffolds, resulting in heterogeneous ability to differentiate (Fig 7c, Fig. S9b). Mouse satellite cells paralleled the behaviors observed with SMPCs cultured onto myoscaffolds, including the ability to differentiate more efficiently on WT and *mdx* ^TG^ samples. Three-dimensional image analysis highlighted the integration and remodeling of satellite cells on WT myoscaffolds, and lack thereof in *mdx* fibrotic regions (Fig. 7d).

To test whether cell behavior on myoscaffolds could be extended to *in vivo* systems, we next transplanted SMPCs into C57-NSG and *mdx*-NSG mice. After 30 days, engrafted tissues were stained for laminin and human lamin A/C to demarcate the engrafted cells (Fig. 7e, Fig. S9c). Similar to the *in vitro* myoscaffolds, laminin intensity widely varied by region in the *mdx* samples. The *mdx* ^NS^ regions exhibited reduced peak laminin intensity compared to C57-NSG, whereas *mdx* ^S^ regions were more than twice as bright (*P<0.05*) (Fig. 7f). As in our *in vitro* assay, we identified SMPCs in *mdx* ^NS^ regions; however, SMPCs were rarely found within *mdx* ^S^ areas as demarcated by dense laminin deposition, unable to infiltrate fibrotic scars. Interestingly, analysis of the basement membrane surrounding engrafted cells revealed similar laminin intensity as that observed globally in WT and *mdx* ^NS^ muscle (Fig. 7g), whereas we saw roughly a 45% reduction in laminin intensity surrounding SMPCs that were able to engraft in *mdx* ^S^ regions, suggesting SMPC integration is dictated by the ability to remodel laminin. These results suggest that incorporation of engrafted cells into the fibrotic scars of dystrophic *mdx* muscle is likely limited by an inability to remodel laminin and that cells may more readily integrate into regions of *mdx* muscle without scarring.

## DISCUSSION

The use of a myoscaffold platform enabled a key finding that cell adhesion is diminished on fibrotic scars characterized by dense laminin deposition, while compensatory ECM production, as observed in *mdx* ^TG^ muscle and myoscaffolds, supports cell adhesion and differentiation. In dystrophic muscle, there is a significant reduction in the availability of laminin receptors due to the loss of dystrophin ^70,71^. We found that laminin disorganization in *mdx* myoscaffolds is heterogeneous and progressive, with reduced laminin deposition in *mdx* ^NS^ myoscaffolds without scar tissue formation, and severe disorganization along with increased laminin deposition with progressive fibrosis (*mdx* ^S^). This disorganization in fibrotic scars is likely reflective of the demands for continual laminin deposition and basement membrane assembly over the repeated cycles of myofiber degeneration and regeneration. As such, we conclude that loss of myofiber adhesion in *mdx* muscle causes disorganization of laminin in the basement membrane that likely renders integrin binding sites inaccessible *in vivo*, and prevents SMPC-integrin receptor binding in the myoscaffold *in vitro* assay. Similarly, we found that SMPCs were unable to remodel the laminin sublayer in fibrotic scars *in vitro* and *in vivo* and displayed impaired differentiation *in vitro*. Sarcospan-induced upregulation of laminin-binding adhesion complexes in *mdx* muscle increases myofiber attachment to laminin, which we hypothesize facilitates normal basement membrane assembly and accessibility of integrin binding sites, which are available for SMPC adhesion. Our results reveal that laminin disorganization is a feature of pathological fibrosis and scarring and presents a barrier for effective muscle regeneration.

We found that the significant laminin α2 deposition in fibrotic scars of *mdx* myoscaffolds was resistant to SMPC-mediated degradation. In fact, the fibrotic scars are impenetrable to SMPCs, inhibiting their growth and differentiation. Degradation of laminin α2 in the basement membrane is an initiating step in myogenesis ^18^. Rayagiri and colleagues demonstrated that, upon activation, satellite cells upregulated production of matrix metalloproteinase (MMP) 2 and 9, permitting digestion of laminin α2 in the basement membrane ^18^. Failure to degrade the basement membrane impairs satellite cell expansion and self-renewal ^18^. Our findings support the premise that fibrotic scars are a major barrier to effective ECM remodeling and that laminin degradation is required for muscle regeneration. Interestingly, we found no significant difference in MMP gene expression levels between SMPCs cultured on WT or *mdx* myoscaffolds.

Although we discovered that laminin disorganization is a primary contributor to impaired stem cell adhesion and differentiation, we observed other changes to the basement membrane, included a reduction of collagen IV and VI, in *mdx* myoscaffolds that could be associated with inhibited myogenesis. Collagen IV is crosslinked to laminin and provides a site of attachment for satellite cells through integrin binding ^72^. A decrease in collagen IV may alter the organization of laminin and reduce the number of binding sites available for satellite cells. Collagen VI has been identified as a key component responsible for regulation of mechanical properties of the satellite cell niche. A decrease in collagen VI abundance has been linked to impaired muscle regeneration and decreased satellite cell self-renewal capacity following injury ^67^. Given the interaction of collagens IV and VI with satellite cells, their decreased abundance in the *mdx* ECM may contribute to failed muscle regeneration in DMD. The observed restoration of collagen VI in the *mdx* ^TG^ ECM may be one mechanism underlying the improved muscle pathology in *mdx* ^TG^ muscle.

Substrate stiffness is known to regulate stem cell differentiation ^73-76^. Fibrosis and increased matrix stiffness cause a myogenic to fibrogenic conversion of stem cells cultured on acellular ECM from aged muscle ^28^. Live cell imaging revealed that SMPCs were unable to deform regions of the *mdx* myoscaffolds containing fibrotic scars. Our AFM data revealed that fibrotic regions are significantly stiffer than both WT controls and regenerating regions of *mdx* myoscaffolds. In fact, cells that settled into the scars during initial cell seeding were unable to migrate and assumed a rounded appearance, indicative of cell stress. We observed an overall mislocalization and disorganization of matrix proteins in the fibrotic *mdx* ECM, indicating disruption of the microenvironment surrounding and separating resident cell populations. Although stem cell adhesion to the basement membrane is a main focus of this report, we recognize that the interstitial matrix provides a niche for many cell types. Given the bidirectional communication between resident cells and their ECM, our findings suggest that the aberrant ECM observed in fibrotic scars could influence the function of as many as ten different mononuclear cell types found in adult skeletal muscle ^77^. Myoscaffolds can be used to investigate the interaction of specific ECM regions with these different cell types, providing further insight into cell-ECM interactions and fate changes in muscle pathology.

The robust remodeling observed in the *mdx* ^TG^ muscle was striking given the significant increase in collagens which are commonly associated with fibrosis and failed regeneration. The increase in collagens III, V, and XI was accompanied by a significant increase in the abundance of collagen crosslinks in *mdx* ^TG^ muscle, most notable in immature crosslinks that are known to confer increased tissue compliance. Therefore, the increased abundance of collagens may be a compensatory strategy to improve force transmission in the *mdx* ^TG^ muscle without increasing ECM stiffness. Further studies are needed to investigate the molecular pathways leading to increased collagen crosslinking and how crosslinking alone affects the ability to remodel the ECM. Interestingly, collagens V and XI are typically observed only during skeletal muscle development. Collagen XI is specifically found only in articular cartilage and intervertebral discs. While the function of collagen XI in skeletal muscle is unknown, Baghdadi and colleagues recently reported that satellite cells produce collagen V that is critical for the calcitonin receptor and notch signaling cascade, which maintains satellite cells in a quiescent state ^78^. Taken together, the changes observed in the *mdx* ^TG^ ECM suggest that upregulation of these developmental and cartilaginous collagens may be beneficial for muscle regeneration. This compensatory ECM phenotype additionally reveals targets that could be leveraged to develop improved therapeutic strategies to address muscle pathology in several disease contexts.

One strategy to improve muscle pathology in DMD is to introduce stem cells that can engraft into skeletal muscle and provide a source of healthy myofibers. In this report, we used SMPCs from human induced pluripotent stem cells to demonstrate the effects of the ECM environment of cell functions necessary for regeneration, particularly since they have promising therapeutic potential. Our findings suggest that modulation of the ECM environment is necessary to enhance the regenerative potential of SMPCs or muscle stem cells, as sarcospan overexpression resulted in improved muscle stem and progenitor cell function in *mdx* myoscaffolds. In further support of this premise, laminin 111, the laminin isoform present in the basement membrane of fetal tissue, was recently shown to have therapeutic potential in dystrophic muscle, leading to improvements in muscle pathology and providing protection from exercise-induced damage following systemic injection ^79^. Upregulation of integrin alpha 7 was observed following laminin 111 treatments, indicating that improved ECM adhesion may underlie the enhanced muscle regeneration. Other known modifiers of DMD muscle pathology, such as osteopontin and LTBP4, are involved in pro-fibrotic signaling, and their reduction diminishes fibrosis and improves muscle pathology ^80-82^. We previously showed that TGF-ß inhibition significantly improves muscle stem cell engraftment ^34^. That LTBP4 sequesters TGF-ß expression further supports how modulation of the ECM environment can be beneficial for SMPC engraftment.

Our study demonstrates the utility of myoscaffolds as a novel tool to study cell-ECM interactions and to test the efficacy of cell-based therapies, building on other elegant systems ^83,84^. Several important biological components of the ECM, including receptor binding sites and mechanical properties, cannot be evaluated or replicated in standard culture dishes or in engineered matrices that lack native architecture and composition. Many different cell types, including fibroblasts and immune cells that are highly active during inflammation, form the skeletal muscle ECM. While we are currently unable to isolate the contribution of each of these cell types to ECM organization, the myoscaffolds can be generated from any tissue, providing insight into ECM deposition and organization in health and disease. Use of the myoscaffold platform allows for direct interrogation of dynamic reciprocity between the ECM and multiple cell types, not only in skeletal muscle, but in different tissues and organ systems. The requirement of small volumes of tissue for analysis makes the application to human biopsies easily translatable. Findings from our use of myoscaffolds additionally inform the creation of engineered matrices designed to investigate the influence of specific matrix components on cell function. Our findings provide proof of concept that the *in vitro* myoscaffold assay is an important tool for direct interrogation of inside-out and outside-in communication.

One limitation of our study is our naïve understanding of the transient changes that occur in the ECM during regeneration. In healthy skeletal muscle, myofibers completely regenerate following injury without any fibrotic scar deposition. Animal models aiming to replicate injury to wild-type muscle often employ cardiotoxin, notexin, or barium chloride injection, accompanied by complete regeneration and virtually no fibrosis within 28 days ^69^. The disease process in *mdx* muscle involves asynchronous cycles of degeneration and regeneration occurring throughout life. In our analysis of *mdx* muscle, we identified ECM regions with myofiber hypertrophy and minimal ECM deposition and other regions with dense fibrotic scars associated with late-stage disease and failed regeneration. In our BaCl2 injury model, we observed an overall increase in SMPC adhesion to myoscaffolds isolated from muscles 7 days post-injury, indicating that there may be inherent properties in the ECM that support improved regeneration early in muscle recovery, despite disease state. While our findings provide interesting insights into the complex cell-ECM interactions shaping ECM biochemical and biophysical properties, they provide only a snapshot into ECM composition during regeneration in health and disease. Further studies are needed to evaluate the chronological expression and organization of ECM proteins in the days and weeks following injury in both healthy and diseased tissues to determine what differences exist that may influence regeneration. Identification of matrix factors that encourage successful myofiber regeneration could provide targets for the development of interventions aiming to improve regeneration and matrix remodeling in early and late stages of muscle disease.

In conclusion, we find that laminin deposition in fibrotic scars presents an impenetrable barrier that impairs cell adhesion and blocks differentiation needed for muscle regeneration. Thus, the loss of cell adhesion to laminin, and not excess collagen deposition, contributes to pathological ECM deposition and distinguishes it from compensatory ECM deposition, which supports effective regeneration. Our work further highlights laminin scarring as a barrier in cell based therapies and suggests engraftments should either be performed prior to the development of extensive fibrotic scarring or that the muscle should be pre-treated with anti-fibrotic agents to reduce the laminin barrier prior to cell transplantations.

## ACKNOWLEDGMENTS

R.H.C. would like to acknowledge early discussions of decellularized skeletal muscle with Dr. Roger P. Farrar (The University of Texas, Austin). The authors would also like to acknowledge Dr. Elizabeth Gibbs for generating the transgenic mice used in the current report and Dr. Jackie McCourt for thoughtful discussion. This work was supported by a Quantitative and Computational Biology (QCB) Collaboratory Postdoctoral Fellowship to Y.Z.K., and the QCB Collaboratory community directed by Dr. Matteo Pellegrini (UCLA). This work was supported by grants from the National Institutes of Health (AR048179 and HL126204 to R.H.C.; AR064327 to A.D.P.; CA046934 to K.C.H.; AR065972 to K.S.R., J.C.R., and M.R.H.; AR052646 to K.S.R.); the UCLA Clinical and Translational Science Institute (UL1TR000124 to R.H.C and K.S.R.); the Muscular Dystrophy Association USA (274143 to R.H.C.); the Broad Stem Cell Research Fellowship, Schaffer Family Foundation Fellowship, and MDA development award (to M.R.H. 629098). We would like to acknowledge the MCDB/Broad Stem Cell Research Center microscopy core at UCLA that provided imaging equipment and support for our investigations, including the Nano and Pico Characterization Laboratory and the Technology Center for Genomics and Bioinformatics. We thank Siobhan McCarthy, Ramiro Diaz-Lara, Daniela Contreras, and Judd Collado for assistance with tissue sectioning, immunofluorescence analysis, and data processing.

## AUTHOR CONTRIBUTIONS

K.S.R. and R.H.C. conceived of and oversaw the project. K.S.R., R.H.C., and K.G.H. developed the on-slide decellularization assay. L.R.S, T.T.P, and K.C.H. performed the proteomic analysis, and K.S.R. and R.H.C. interpreted the data. K.S.R. initiated the AFM experiments, K.S.R., A.S., N.A.G., S.H., and S.K. designed the experiments, K.S.R. and S.K. performed the experiments, and K.S.R. and S.H. analyzed the data. K.S.R. designed the ECM component localization experiments, K.S.R. and K.H. performed the experiments, A.M. created the MATLAB code, A.M., K.G.H., and J.C. processed the data and K.S.R. analyzed the results. M.R.H., K.S.R., and K.Y. designed the live cell imaging experiments, K.Y. performed the imaging, and K.S.R., M.R.H., and K.Y. analyzed the data. M.R.H. generated the cell lines used in all the experiments and K.S.R. and M.R.H. designed and conducted the cell adhesion, proliferation, and differentiation experiments. Y.Z.K. analyzed the RNA sequencing data and K.S.R. and M.R.H. interpreted the results. M.R.H. generated the NSG mice and performed the iPSC engraftment experiments and M.R.H. and K.S.R. performed the tissue analysis. K.S.R. and R.H.C. created the remodeling index and K.S.R. quantified ECM remodeling *in vivo* and *in vitro*. K.S.R., J.C.R., and R.H.C. designed the barium chloride injury experiments. K.S.R. and J.C.R. performed the experiments, processed the data, and analyzed the results. K.S.R. and R.H.C. wrote the manuscript with contributions from M.R.H. on analysis and interpretation of results from the cell studies. A.D.P., B.G.N., K.C.H., and R.W. contributed to discussion of results and edited the manuscript. R.H.C., A.D.P., B.G.N., K.C.H., A.S., N.A.G., S.K., and R.W. provided resources for the investigation.

## MATERIALS AND METHODS

### Study Design

The objective of the current study was to utilize a reductionist approach to investigate the effects of the skeletal muscle ECM in DMD on specific functions of stem cells that are necessary for regeneration and, reciprocally, to determine how cells interact with and modulate the ECM in the absence of dystrophin. We utilized the *mdx* mouse as the murine model of DMD, and generated the *mdx* ^TG^ mice to investigate modulation of laminin organization and deposition in the DMD ECM. Investigators were not blinded to the experimental groups. All experiments were performed using n-values based on a priori power analysis calculations and as determined by previous experience. The numbers of animals, cells, and biological replicates are indicated in the figure legends. All mice used in the study were maintained in the Terasaki Life Sciences Vivarium following guidelines established by the Institutional Animal Care and Use Committee at the University of California, Los Angeles (approval #2000-029-43) and approval for these studies were granted by the UCLA Animal Welfare Assurance (approval #A3196-01). All human pluripotent stem cell (hPSC) work was approved by ESCRO. Experiments were performed using skeletal muscle progenitor cells derived from three different wildtype hPSC lines and one murine satellite cell line (see below for specific cell line generation protocols). Cell lines used for each experiment are specified in the figure legends.

### Animals

Wild-type (C57BL/6J) and *mdx* mice were purchased from Jackson Laboratories (Bar Harbor, ME, USA). To generate the *mdx*:SSPN-Tg (*mdx* ^TG^) mice, we first generated human SSPN (hSSPN) transgenic mice expressing full-length hSSPN cDNA under control of the human skeletal α-actin promoter. Transgenic constructs were designed with a SV40 VP1 intron located downstream of the human skeletal α-actin promoter, as described previously ^25,59,85^. Male SSPN-Tg mice were then crossed to *mdx* females to generate dystrophin-deficient mice expressing transgenic hSSPN, as previously described ^85^.

All mice used in the study were 18-22 week old males, except for those used in the BaCl2 injury experiments, which were 11-16 weeks of age. For all experiments, except mass spectrometry and BaCl2 injury experiments, the quadriceps muscle was dissected and mounted in OCT (Tissue-Tek, Sakura Finetek, Torrance, CA, USA) and flash frozen in liquid nitrogen–cooled isopentane. Tissues were then stored at −80°C until further processing. For proteomic analysis and BaCl2 injury experiments, the quadriceps muscle was snap frozen in liquid nitrogen and stored at −80°C.

### Cell Lines

All human pluripotent stem cell (hPSC) work was approved by ESCRO. Throughout the study, we performed experiments on skeletal muscle progenitor cells derived from three different wildtype hPSC lines. We used one human embryonic stem cell line H9, (obtained from Wicell) and two human induced pluripotent stem cell lines. Fibroblasts taken from patient skin biopsies at the Center for Duchenne Muscular Dystrophy (CDMD) were reprogrammed to derive 1002 (wild-type) hiPSC lines as previously described ^86^. For live cell imaging, the hiPSC line WTC-11 (derivation reported in Kreitzer FR, et al.^35^, obtained from the Allen Institute) contained an mTagRFP transgene inserted in the AAVS1 safe harbor locus, which constitutively expresses red fluorescence at the plasma membrane. HPSCs were grown and maintained on hESC-Qualified matrigel-coated plates in mTESR medium (Stem Cell Technologies) containing 0.4% P/S (Hyclone). Directed differentiation of hiPSCs to skeletal muscle progenitor cells (SMPCs) was performed as previously described ^34^. After 50 days of differentiation, SMPCs were enriched using HNK1-ERBB3+NGFR+ surface markers by flow cytometry. Enriched hPSC-SMPCs were maintained in SkBM2 (Lonza) for 3-7 days prior to seeding onto decellularized ECM.

Murine satellite cells used to investigate cell adhesion and differentiation were isolated from a Pax7-Cre Rosa26-ZsGreen mouse model that constitutively fluoresces after tamoxifen induced Cre-recombination. To isolate satellite cells, we pooled the tibialis anterior, gastrocnemius, and quadriceps muscles, bilaterally from N=6 male C57Bl/10 Pax7-Cre ZsGreen mice (10 weeks old). Satellite cells were sorted (Cd45-Ter119-ZsGreen+) and immediately seeded on WT, *mdx*, and *mdx* ^TG^ myoscaffolds for analysis.

### Muscle Decellularization

In the development of our decellularization protocol, we considered and refined many variables that influence the production of acellular scaffolds including tissue section thickness and mounting material required to prevent sample loss during the decellularization procedure (data not shown). Based on extensive testing, we found that 30-50μm sections mounted on slides with an adhesive coating (#FF-914 Matsunami Glass Ind. Ltd., Kishiwada, Osaka, Japan) were optimal for retaining sections during decellularization.

Transverse cryosections from the quadriceps muscle of WT, *mdx*, and *mdx* ^TG^ mice (30 μm for H&E and IFA, 50 μm for AFM and cell seeding studies) were placed onto adhesive slides (#FF-914 Matsunami Glass Ind. Ltd., Kishiwada, Osaka, Japan) and allowed to dry at room temperature for 2 hours. For sections designated for atomic force microscopy, 10 uL of Tissue Tack (Electron Microscopy Sciences, Hatfield, PA, USA) was applied to the slide prior to section placement. Following drying, slides were placed in 1% SDS and decellularized at room temperature under constant rotation (50 rpm) for 10-60 minutes. Slides were then placed in 50 mL 1x PBS for 30 minutes, followed by 50 mL diH_2_0 for 30 minutes, and ending with a final rinse in 50 mL of 1x PBS for 30 minutes. Decellularized sections were then stored in 1x PBS and used on the same day they were produced.

### Histology

To assess general muscle pathology and the removal of cellular material, untreated and decellularized sections were stained with hematoxylin and eosin, as described previously ^58^. Images were captured under identical conditions using an Axioplan 2 fluorescent microscope equipped with a 10x and 20x differential interference contrast objectives and the Axiovision Rel 3.0 software (Carl Zeiss, Inc., Thornwood, NY, USA).

### Immunohistochemistry

Decellularized and untreated muscle sections were blocked with 3% BSA in PBS for 30 min at room temperature. Avidin/biotin blocking kit (Vector Laboratories) was used according to manufacturer’s instructions. For antibodies raised in mouse, Mouse on Mouse blocking reagent (Vector Laboratories) was used according to manufacturer’s instructions. Sections were incubated in primary antibody in PBS at 4°C overnight with the following antibodies: collagen I (CL50151AP-1; 1:250; Cedarlane Labs), collagen IV (AB19808; 1:250; Abcam), collagen VI (70R-CR009X; 1:200; Fitzgerald Industries), fibronectin (AB2413; 1:250; Abcam), laminin (L9393; 1:500; Sigma-Aldrich) and laminin alpha-2 (AB11576; 1:150; Abcam). Primary antibodies were detected by biotinylated anti-rabbit (BA-1000; 1:500; Vector Laboratories). Fluorescein-conjugated avidin D (A-2001; 1:500; Vector Laboratories) was used to detect secondary antibodies. Laminin alpha-2 was used specifically for costaining and was detected by Alexa Fluor 594 Donkey anti-rat (AB150156; 1:200; Abcam). All sections were mounted in Vectashield (Vector Laboratories) and visualized using either an Axioplan 2 fluorescence microscope with Axiovision 3.0 software (Carl Zeiss Inc., Thornwood, NY, USA) or a Leica TCS SP5 confocal microscope equipped with an argon 488 nm and helium-neon 594 nm. For confocal microscopy, images were taken at 1024 x 1024 resolution with 20X and 63X oil objectives using LAS X software.

### Cell Proliferation and Differentiation Assays

Transverse cryosections (50 μm) from the quadriceps of WT, *mdx*, and *mdx* ^TG^ mice were placed on MAS slides (Matsunami Glass Ind. Ltd., Kishiwada, Osaka, Japan) and arranged to cover the whole surface of the slide. Slides were allowed to dry at room temperature for 2 hours and stored at −80°C. Prior to decellularization, chambers from Lab-Tek Chamber slides (Nunc, Rochester, NY, USA) were removed and placed over the MAS slide, ensuring that the borders of the chamber were not in contact with any muscle sections. The chambers were then sealed with silicone (Flexbar reprorubber, Islandia, NY, USA) and placed in a chemical fume hood to dry for one hour. 1% SDS was then pipetted into each chamber and slides were placed on an orbital rotator (50 rpm) for 40 minutes. The SDS was removed, 3 mL of PBS was added to each chamber, and the slides were again placed on the rotator for 30 minutes. Three more rinses, one in ddH20, followed by two in PBS, were performed, for 30 minutes each, all under constant rotation. Following the final rinse, the PBS was removed and 3 mL of SkBM-2 (Lonza) media was added to each chamber. Slides were then placed in an incubator at 37°C and left overnight.

The following day, ERBB3+NGFR+ hPSC SMPCs generated from H9, CDMD 1002, and WTC-11 lines and cultured in 6-well plates were dissociated with TRPYLE (Stem Cell Technology), centrifuged at 300 g, and pelleted. SMPCs were resuspended in SkBM-2 (Lonza) and 150,000 cells per chamber slide were seeded. Slides containing myoscaffolds without cells were maintained alongside those with cells and served as controls for all experiments. Myoscaffolds with and without HPSC SMPCs were maintained in SkBM-2 proliferation media for 5 days, or until they reached 90% confluence. For RNA sequencing experiments, forceps were used to remove matrices and attached SMPCs. The ECM and attached cells were immediately placed in lysis buffer and RNA extracted using RNEasy minikits (Qiagen) for RNA-Sequencing. Three to four matrix sections were left on each slide and then fixed in 4% PFA for immunostaining. Immunostaining of collagen, laminin, and phalloidin were performed as previously described ^87^.

For live cell imaging, WTC-11 hiPSCs containing an mTagRFP transgene inserted in the AAVS1 safe harbor locus, which constitutively expresses red fluorescence at the plasma membrane, were obtained from the Coriell Institute. HiPSCs were differentiated to SMPCs, enriched for ERBB3+NGcV VZXadFR+, and maintained in SkBM-2. Acellular ECM from WT, *mdx*, and *mdx* ^TG^ mice were stained using pan-Laminin and taken to the Zeiss Spinning Disk Confocal for imaging (UCLA MCDB / BSCRC microscopy core). Z-stack images were set based on laminin fluorescence. Immediately upon cell seeding, cells were imaged in 10-minute intervals over 4 days.

To induce myotube differentiation, SMPCs were switched to N2 media (containing 1% N-2 supplement (Thermo Fisher), 1% Insulin Transferrin Selenium, 10ng/ml IGF-1, and 3μM SB-431542 in DMEM/F12) for 5 days. HPSC myotubes on acellular matrix were fixed in 4% PFA and immunostained for α-actinin (Sigma) or myosin heavy chain (MYH1E; MF-20; 1:20; DSHB) and DAPI. To measure cell-mediated basement membrane remodeling of acellular matrix, chamber slides were additionally stained for Laminin-α2. Images of myotubes were taken from six-eight independent WT, *mdx*, and *mdx* ^TG^ cross-sections for quantification. Fusion efficiency of myotubes was calculated by counting DAPI+ nuclei contained in α-actinin+ myotubes containing 3 or more nuclei, as a percentage of all DAPI nuclei within the field of view. Cells that were α-actinin + containing one to two DAPI+ nuclei were assigned as myocytes, and not included in myotube fusion index.

To demonstrate cellular integration within the matrix, 20X and 63X images were acquired using the Zeiss LSM780 confocal microscope. Optical sections were imaged using Zen Blue 2.1 optimal recommended settings. Images of *mdx* ^NS^ and *mdx* ^S^ myoscaffolds were selected based on Laminin-α2 density and encroachment into the interstitial space. To visualize myotubes in X-Z plane, images were transferred to Imaris 9.2 (Bitplane) and surface rending applied to chromatic channels. Images were rotated and the Clippling Plane feature used to view the myotube sagittal plane. Imaris Vantage feature was used to calculate the Intensity mean location of α-actinin+ cells in the Z plane.

### ECM Crosslinking

Approximately, 5mg of tissue was reduced with NaBH_4_, then hydrolyzed with 6N HCl. Acid was removed under vacuum and the samples were enriched using cellulose resin. Samples were dried under vacuum and resuspended in 0.1% FA for LC-MS analysis. Crosslinks were detected with a QExactive mass spectrometer (Thermo Fisher Scientific, San Jose, CA, USA) coupled with a Vanquish UPHLC system (ThermoFisher, San Jose, CA, USA). Chromatographic separation was achieved using an Acquity UHPLC BEH Amide column (2.1 x 100mm, 1.7μm particle size – Waters, Milford, MA, USA), and data was acquired using a full MS scan in positive polarity. Accurate mass, retention time matching to standards, and MS2 validation were used for peak assignment. Data was converted to mzXML file format and analyzed using El-MAVEN (Princeton, NJ, USA).

### Atomic Force Microscopy (AFM)

AFM measurements were performed on 50 μm decellularized muscle sections using an MFP-3D_BIO system (Asylum Research, Oxford Instruments, Goleta, CA, USA). The endomysium of each sample was probed with Olympus 200 µm long TR400PB probes (nominal freq (kHz)= 32(20-49), k(N/m)= 0.02(0.01-0.05); Asylum Research, Oxford Instruments, Goleta, CA, USA). The sensitivity and spring constant of each probe were calibrated before each experiment using the automated GetReal routine. All measurements were taken in contact mode and force-vs-indentation curves were generated from an average of 100 points/sample. Approach and retraction speeds for all trials were 2 μm/sec with a trigger force of 3 nN. Data analysis was performed using Asylum Research software (Version 16). To evaluate ECM stiffness, Young’s modulus was calculated for each curve, using the Hertz-Sneddon model ^88^. The overall adhesive properties of the tissue were determined as the peak rupture force during the retraction of the AFM probe away from the test sample surface ^89^. The average Young’s modulus and adhesive force were then calculated and reported for each sample.

### Cell Adhesion Assays

HPSC SMPC’s (CDMD 1002 and H9 cell lines) were seeded onto myoscaffolds at density of 50,000 cells/3 sections and placed in an incubator. After 4 hours, media was removed and slides were rinsed in 1x PBS for 30 sec. Slides were then treated with Gentle dissociation buffer (Stem Cell Technologies) and placed on an orbital rotator for 15 minutes @ 50rpm (room temp). Non-dissociated slides served as a pre-dissociation control. All slides were then washed in PBS and fixed in 4% PFA for 10 minutes, followed by 2 rinses in 1x PBS for 10 seconds.

Primary mouse satellite cells required longer to adhere and were cultured for 16 hours before dissociation. Satellite cells were more adherent to myoscaffolds relative to hPSCs SMPCs, thus we increased the EDTA concentration (2mM), NaCl osmolality (250mM), and time in buffer (20 mins) to increase stringency of the assay. As primary mouse satellite cells constitutively expressed a ZsGreen tag this enabled us to measure the same tissue regions pre and post-dissociations without the need for additional staining.

For cell adhesion assays using SMPCs, slides were stained with phalloidin and DAPI for identification of cell number pre- and post-dissociation. Immunostaining of laminin was also performed as previously described and was utilized to distinguish the ECM boundaries for determination of cells on the myoscaffold relative to those on the glass slide ^87^.

### RNA-Sequencing

RNA isolated from hPSC SMPCs cultured on acellular matrices were taken to the UCLA Technology Center for Genomics and Bioinformatics for analysis. Libraries for RNA-Seq were prepared with KAPA Stranded mRNA-Seq Kit. The workflow consists of mRNA enrichment, cDNA generation, and end repair to generate blunt ends, A-tailing, adaptor ligation, and PCR amplification. Different adaptors were used for multiplexing samples in one lane. Sequencing was performed on Illumina HiSeq 3000 for 1×50 run. Data quality check was done on Illumina SAV. Demultiplexing was performed with Illumina Bcl2fastq2 v 2.17 program. Total numbers of sequenced reads were 17-20 million per sample. RNA-Seq reads were mapped to the human reference genome (hg38) using STAR aligner ^90^. For each sample, 15-18 million reads were uniquely mapped to the genome (88% of sequenced reads) and were used to measure expression of all annotated genes (Ensembl v.92). Raw gene counts were normalized to CPM (counts-per-million). Lowly expressed genes were removed, and 9831 genes were kept for further analysis (with CPM > 10 in at least 2 samples). Differential expression analysis was performed using EdgeR/LRT ^91^. Differentially expressed genes (DEGs) were identified at a false discovery rate of 5% (FDR, q<0.05) and a minimum 1.5 log fold-change was used as a cut-off.

### qPCR

As for RNA-Seq experiments, RNA was immediately isolated from SPMCs following cell seeding experiments using RNeasy Microkits (Qiagen). Primers for CFOS, LAMA2, ITGA7 and GAPDH were designed using NCBI primer blast or based on published work (CFOS: forward sequence = GTTGCCACCCCGGAGTCTGAG, reverse sequence = GCCTGGATGATGCTGGGAACA; LAMA2: forward sequence = GGAACTACCCTCGCTGCAAT, reverse sequence = GGCATCGAGTCCGAATTT; ITGA7: forward sequence = AATCTGGACGTGATGGGTGC, reverse sequence = TCAGTCTCCTCCAGGCTCAA; GAPDH: forward sequence = CGCCCCCGGTTTCTATAAATTG, reverse sequence = AAGAAGATGCGGCTGACTGT). Primers (10 µm) were validated using cells known to express the genes described. Complementary DNA (cDNA) concentrations were 5-fold serially diluted starting at 5 ng/µl. Primers with 0.9–1.1 efficiency were used for experiments.

### Proteomics

Quadriceps muscles were harvested from WT, *mdx*, and *mdx* ^TG^ mice and snap frozen in liquid nitrogen. Samples were then prepared for mass spectrometry analysis as previously described ^55^. In short, samples were pulverized in liquid nitrogen using a ceramic mortar and pestle and lyophilized. For each sample, 5 mg (dry weight) of tissue was homogenized in 200mL/mg high-salt buffer (HS buffer) containing a 1x protease inhibitor. Pellets were subjected to three rounds of HS buffer wash followed by treatment with 6M guanidine extraction buffer. The remaining pellets from each tissue, representing insoluble ECM proteins, were digested with freshly prepared hydroxylamine buffer as previously described (ref 52). 100µL of the cellular fraction (combined fractions 1, 2, and 3) and 200µL of the soluble and insoluble ECM fractions were subjected to enzymatic digestion with trypsin using a filter-aided sample prep (FASP) approach and C18 tip cleanup ^92^. Samples were then analyzed by liquid chromatography – data dependent acquisition (DDA) tandem mass spectrometry (LC-MS/MS) as previously described ^93^. Samples were analyzed on a Q Exactive HF Orbitrap mass spectrometer (Thermo Fisher Scientific) coupled to an EASY-nanoLC 1000 system through a nanoelectrospray source. The analytical column (100 μm i.d. × 150 mm fused silica capillary packed in house with 4 μm 80 Å Synergi Hydro C18 resin (Phenomenex; Torrance, CA)) was then switched on-line at 600 nL/min for 10 min to load the sample. The flow rate was adjusted to 400 nL/min, and peptides were separated over a 120-min linear gradient of 2–40% ACN with 0.1% FA. Data acquisition was performed using the instrument supplied Xcalibur (Thermo Fisher Scientific, San Jose, Calif) software in positive ion mode. MS/MS spectra were extracted from raw data files and converted into mgf files using a PAVA script (UCSF, MSF, San Francisco, Calif). These mgf files were then independently searched against mouse SwissProt database using an in-house Mascot server (version 2.2.06; Matrix Science, London, UK). Mass tolerances were +/−10 ppm for MS peaks and +/−0.5 D for MS/MS fragment ions. Trypsin specificity was used allowing for one missed cleavage. Pro oxidation (hydroxy proline, indicative of hydroxylation, delta mass = 15.9949), Met oxidation (methionine, delta mass = 15.9949), protein N-terminal acetylation (delta mass = 42.0106), and peptide N-terminal pyroglutamic acid formation (delta mass = −17.0265) were allowed for variable modifications, whereas carbamidomethyl of Cys (delta mass = 57.02146) was set as a fixed modification.

Following Mascot searches, data was directly loaded into Scaffold^TM^ (Proteome Software Inc.). Peptide spectral matches (PSMs) were directly exported with a 99% confidence in protein identifications and at least 2 unique peptides per protein, resulting in a false discovery rate of 0.54%. 2-group comparisons were done by two-sided Student’s t-tests followed by Bonferroni correction to account for multiple comparisons. Partial least squares-discriminant analysis (PLSDA) was performed using MetaboAnalyst (version 3.0) with sum and range scaling normalizations ^94^.

### Quantification of ECM Component Abundance and Localization

Image quantification was performed on confocal images (63X) using Image J software (NIH, version 1.50i) for each of the following co-stains pairs: laminin alpha-2 and collagen I, laminin alpha-2 and collagen IV, laminin alpha-2 and collagen VI, and laminin alpha-2 and fibronectin. Image J was used to merge individual red (laminin alpha-2) and green (collagens, fibronectin) channels to create a composite channel overlay. Using the line drawing tool to quantify regions of interest, lines were drawn perpendicularly to the endomysium previously bordering two muscle fibers (Fig. S4b). Areas that lacked a distinct basement membrane were not considered for analysis. A plot profile was created for each red and green channel and data was exported for compilation in excel.

Excel files were processed in MATLAB (The MathWorks, Inc., R2018b (Version 9.5)) using a custom algorithm. The code first calls microscopic red and green channels as raw data and scales the data with respect to its basal value. The data are then filtered using MATLAB filtfilt function (filtfilt(d,x) zero-phase filters of the input data, x, using a digital filter, d). Using data first from the red channel, the code finds the two maximum peaks and their locations. Each peak belongs to a distribution and it is fitted with MATLAB multiple (gauss2) or single (gauss1) Gaussian function depending upon the nature of distribution. The program optimally sets the minimum and maximum cut off from a fitted distribution along the X-axis considering two standard deviations from the mean to account for 95% area within. Four segment points were quantified for each red channel and these points were then applied to the corresponding green channel data. The area under the curve was calculated to represent the abundance of each component in the respective regions. For each ECM protein component, data were compiled in GraphPad Prism (Version 8.4.2 (464)) and outliers were removed using the ROUT method.

### Barium Chloride Injury Model and Myoscaffold Preparation

To generate the BaCl_2_ injury model for analysis of SMPC function on regenerating myoscaffolds, the tibialis anterior (TA) of WT, *mdx*, and *mdx* ^TG^ mice was injected with 50 μL of 1.2% BaCl_2_ dissolved in PBS ^69^. Prior to injections, mice were anesthetized using 2% isofluorane. Tissues were collected 4 days and 7 days after BaCl_2_ injection, snap frozen in liquid nitrogen, and stored at −80°C until further processing.

To generate myoscaffolds, transverse cryosections (25 μm, sectioned at −20°C) from the tibialis anterior muscle of WT, *mdx*, and *mdx*^TG^ mice (7 days post-injury) were placed on adhesive slides (#FF-914 Matsunami Glass Ind. Ltd., Kashiwada, Osaka, Japan) and allowed to dry at room temperature for two hours. Once dry, slides were placed in 1% SDS and decellularized at room temperature under constant rotation (50 rpm). Given that the TA muscle was undergoing active regeneration at 7 days, we conducted additional decellularization experiments and found 10 minutes to be the optimal exposure time in 1% SDS to retain ECM composition and remove cellular material from BaCl_2_ treated muscles (Fig. S8c). Following SDS treatment, slides were then placed in 50 mL 1x PBS for 15 minutes immediately followed by another 1x PBS wash for 45 minutes, followed by 50 mL diH_2_O for 30 minutes, and ending with a final rinse in 50 mL of 1x PBS for 45 minutes. Decellularized tissues were stored in 1x PBS at 4°C until further use in cell adhesion and differentiation experiments, as described above.

### *In vivo* SMPC engraftment

To generate immunocompromised mice for human muscle cell engraftment, Nod-Scid-Gamma (NSG) mice (Jax #005557) were crossed to C57BL/10J (Jax #00665) or *mdx* (Jax #001801). C57-NSG, *mdx*-NSG were then backcrossed for 5-8 generations to the parental non-NSG strain and genotyped for retention of Scid and IL-2 knockout alleles. *mdx*-NSG mice were housed in the Humanized Mouse Core at UCLA, an immunocompromised core facility. All animal work was conducted under protocols approved by the UCLA Animal Research Committee in the Office of Animal Research Oversight. Congenic C57-NSG and *mdx*-NSG mice were then used for engraftment studies.

After 6-7 weeks of directed differentiation, hPSC-SMPCs were removed from culture using TRIPLE and FACS sorted using HNK1-ERBB3+NGFR+ cell surface markers. SMPCs were then cultured in SkBM-2+bFGF for four to eight days depending on the experiment. SMPCs were then dissociated pelleted at 300g, and resuspended in Hank’s Balanced Salt Solution (HBSS) at 1×10^6^ cells/per 5µls. This allowed us to obtain at least one million cells per mouse for engraftment. To induce muscle damage, mice were pretreated with 1.2% BaCl_2_ 24 hours prior to transplantation. Mice were then anesthetized using 2% isofluorane, and 5-10µl of cells in solution were injected into the tibialis anterior (TA) muscle using Hamilton micro-syringes. A total of three separate directed differentiations were performed using H9 cells and mice from C57-NSG and *mdx*-NSG strains were engrafted from each experiment (N=6 total mice, 3 per strain).

After 30 days, C57-NSG and *mdx*-NSG mice were euthanized and TA muscles dissected. The TA muscles were immediately embedded in optimum cutting temperature (OCT) compound and flash-frozen in isopentane cooled by liquid Nitrogen. Embedded muscles were stored at −80°C until sectioned in 10µm slices using a Leica microcryotome.

To quantify ECM remodeling post cell engraftment, tissues were stained for the human specific marker LaminAC^+^ and a pan-Laminin antibody to demarcate regions of fibrosis or regeneration. We identified regions containing engrafted human cells and further categorized fibrotic vs. regenerative regions in *mdx-NSG* samples based on central nucleation and laminin deposition. Laminin intensity was quantified in the basement membrane directly surrounding human cells (local remodeling) or in regions distant from the human cells (global remodeling). For global laminin intensity, 30 random regions were selected per image, and all measurements were performed using the line drawing tool in Image J (NIH, version 1.50i), as described above.

### Statistical Analysis

All statistical analyses were performed using GraphPad Prism (Version 8.4.2 (464)). For all experiments, comparisons between two groups were performed by two-tailed unpaired t-tests, and comparisons of multiple groups were performed by one-way ANOVA. All n-values and statical analyses are indicated in the respective figure legends. Results are expressed as mean ± SD or SEM, as indicated. P < 0.05 was considered statistically significant in all cases, except for the proteomic analysis where Bonferroni correction was applied to adjust for multiple comparisons.

## DATA AVAILABILITY

All data are available in the main text or the supplementary materials. The complete RNA sequencing data set generated for SMPCs cultured on WT and *mdx* myoscaffolds (Fig. S2c) is available in Supplemental Table 1. The mass spectrometric dataset used for the analysis in Fig. 4a,b and Fig. S3a-c is available in Supplemental Table 2. Any additional data that support the findings of this study are available from the corresponding author upon reasonable request.

## CODE AVAILABILITY

The MATLAB code utilized to determine ECM component distribution is available from the corresponding author upon reasonable request.

---

**Supplemental Table 1: RNA-Seq dataset reveals that *mdx* myoscaffolds affect gene expression differently than wild-type controls.**

List of differentially expressed genes detected in SMPCs (H9 cells) cultured on wild-type (WT, n=3) and *mdx* (n=2) myoscaffolds, including log2-fold-change and *p*-values.

**Supplemental Table 2: Proteomics data**

List of proteins detected in wild-type (WT), *mdx*, and *mdx* ^TG^ quadricep muscle samples (n=5 per group) from mass spectrometry.

**Supplemental Video 1: SMPCs migrate and remodel laminin in WT myoscaffolds.**

Video of SMPCs (WTC-11 cells, red) on a wild-type (WT) myoscaffold (laminin, green) during the first 14 ½ hours (captured in 10-minute increments) following addition of cells. The arrows indicate areas of ECM deformation and laminin remodeling, the latter indicated by a reduction in green fluorescence.

**Supplemental Video 2: SMPCs migrate and circle the basement membrane on a *mdx* ^NS^ myoscaffold.**

Video of SMPCs (WTC-11 cells, red) on a *mdx* ^NS^ myoscaffold (laminin, green) during the first 14 ½ hours (captured in 10-minute increments) following addition of cells. The arrow indicates a cell circling the basement membrane (between 6hr:30sec-10hr:00sec).

**Supplemental Video 3: SMPCs avoid fibrotic scars in *mdx* ^S^ myoscaffolds.**

Video of SMPCs (WTC-11 cells, red) on a *mdx* ^S^ myoscaffold (laminin, green) during the first 14 ½ hours (captured in 10-minute increments) following addition of cells. A large fibrotic region is present in the center of the field of view.

**Supplemental Video 4: SMPCs rapidly circle the basement membrane of *mdx* scaffolds.**

Video of SMPCs (WTC-11 cells, red) on an *mdx* myoscaffold (laminin, green) on the second day following addition of cells. The arrows indicate cells rapidly circling the basement membrane.

## REFERENCES

1 Dumont, N. A., Bentzinger, C. F., Sincennes, M. C. & Rudnicki, M. A. Satellite Cells and Skeletal Muscle Regeneration. Compr Physiol 5, 1027–1059, doi:10.1002/cphy.c140068 (2015).

2 Feige, P., Brun, C. E., Ritso, M. & Rudnicki, M. A. Orienting Muscle Stem Cells for Regeneration in Homeostasis, Aging, and Disease. Cell Stem Cell 23, 653–664, doi:10.1016/j.stem.2018.10.006 (2018).

3 Mauro, A. & Adams, W. R. The structure of the sarcolemma of the frog skeletal muscle fiber. The Journal of biophysical and biochemical cytology 10(4)Suppl, 177-185 (1961).

4 Gumucio, J. P., Sugg, K. B. & Mendias, C. L. TGF-beta superfamily signaling in muscle and tendon adaptation to resistance exercise. Exerc Sport Sci Rev 43, 93–99, doi:10.1249/JES.0000000000000041 (2015).

5 Ervasti, J. M. & Campbell, K. P. Membrane organization of the dystrophin-glycoprotein complex. Cell 66, 1121–1131 (1991).

6 Ervasti, J. M. & Campbell, K. P. A role for the dystrophin-glycoprotein complex as a transmembrane linker between laminin and actin. The Journal of cell biology 122, 809–823 (1993).

7 Ibraghimov-Beskrovnaya, O. et al. Primary structure of dystrophin-associated glycoproteins linking dystrophin to the extracellular matrix. Nature 355, 696–702, doi:10.1038/355696a0 (1992).

8 Pegoraro, E. et al. SPP1 genotype is a determinant of disease severity in Duchenne muscular dystrophy. Neurology 76, 219–226, doi:10.1212/WNL.0b013e318207afeb (2011).

9 Flanigan, K. M. et al. LTBP4 genotype predicts age of ambulatory loss in Duchenne muscular dystrophy. Annals of neurology 73, 481–488, doi:10.1002/ana.23819 (2013).

10 Kramerova, I. et al. Spp1 (osteopontin) promotes TGFbeta processing in fibroblasts of dystrophin deficient muscles through matrix metalloproteinases. Human molecular genetics, doi:10.1093/hmg/ddz181 (2019).

11 Vetrone, S. A. et al. Osteopontin promotes fibrosis in dystrophic mouse muscle by modulating immune cell subsets and intramuscular TGF-beta. The Journal of clinical investigation 119, 1583–1594, doi:10.1172/JCI37662 (2009).

12 Edwards, D. R. et al. Transforming growth factor beta modulates the expression of collagenase and metalloproteinase inhibitor. EMBO J 6, 1899–1904 (1987).

13 Lamar, K. M., Miller, T., Dellefave-Castillo, L. & McNally, E. M. Genotype-Specific Interaction of Latent TGFbeta Binding Protein 4 with TGFbeta. PloS one 11, e0150358, doi:10.1371/journal.pone.0150358 (2016).

14 Peter, A. K., Cheng, H., Ross, R. S., Knowlton, K. U. & Chen, J. The costamere bridges sarcomeres to the sarcolemma in striated muscle. Prog Pediatr Cardiol 31, 83–88, doi:10.1016/j.ppedcard.2011.02.003 (2011).

15 Colognato, H., Winkelmann, D. A. & Yurchenco, P. D. Laminin polymerization induces a receptor-cytoskeleton network. The Journal of cell biology 145, 619–631, doi:10.1083/jcb.145.3.619 (1999).

16 Henry, M. D. & Campbell, K. P. A role for dystroglycan in basement membrane assembly. Cell 95, 859–870, doi:10.1016/s0092-8674(00)81708-0 (1998).

17 Jacobson, C., Cote, P. D., Rossi, S. G., Rotundo, R. L. & Carbonetto, S. The dystroglycan complex is necessary for stabilization of acetylcholine receptor clusters at neuromuscular junctions and formation of the synaptic basement membrane. The Journal of cell biology 152, 435–450, doi:10.1083/jcb.152.3.435 (2001).

18 Rayagiri, S. S. et al. Basal lamina remodeling at the skeletal muscle stem cell niche mediates stem cell self-renewal. Nat Commun 9, 1075, doi:10.1038/s41467-018-03425-3 (2018).

19 Goody, M. F. et al. NAD+ biosynthesis ameliorates a zebrafish model of muscular dystrophy. PLoS Biol 10, e1001409, doi:10.1371/journal.pbio.1001409 (2012).

20 Costa, A., Naranjo, J. D., Londono, R. & Badylak, S. F. Biologic Scaffolds. Cold Spring Harb Perspect Med 7, doi:10.1101/cshperspect.a025676 (2017).

21 Londono, R. & Badylak, S. F. Biologic scaffolds for regenerative medicine: mechanisms of in vivo remodeling. Annals of biomedical engineering 43, 577–592, doi:10.1007/s10439-014-1103-8 (2015).

22 Wolf, M. T., Dearth, C. L., Sonnenberg, S. B., Loboa, E. G. & Badylak, S. F. Naturally derived and synthetic scaffolds for skeletal muscle reconstruction. Advanced drug delivery reviews, doi:10.1016/j.addr.2014.08.011 (2014).

23 Merritt, E. K. et al. Functional assessment of skeletal muscle regeneration utilizing homologous extracellular matrix as scaffolding. Tissue engineering. Part A 16, 1395–1405, doi:10.1089/ten.TEA.2009.0226 (2010).

24 Marshall, J. L., Kwok, Y., McMorran, B. J., Baum, L. G. & Crosbie-Watson, R. H. The potential of sarcospan in adhesion complex replacement therapeutics for the treatment of muscular dystrophy. The FEBS journal 280, 4210–4229, doi:10.1111/febs.12295 (2013).

25 Peter, A. K., Marshall, J. L. & Crosbie, R. H. Sarcospan reduces dystrophic pathology: stabilization of the utrophin-glycoprotein complex. J Cell Biol 183, 419–427, doi:10.1083/jcb.200808027 (2008).

26 Gillies, A. R., Smith, L. R., Lieber, R. L. & Varghese, S. Method for decellularizing skeletal muscle without detergents or proteolytic enzymes. Tissue engineering. Part C, Methods 17, 383–389, doi:10.1089/ten.TEC.2010.0438 (2011).

27 Perniconi, B. et al. The pro-myogenic environment provided by whole organ scale acellular scaffolds from skeletal muscle. Biomaterials 32, 7870–7882, doi:10.1016/j.biomaterials.2011.07.016 (2011).

28 Stearns-Reider, K. M. et al. Aging of the skeletal muscle extracellular matrix drives a stem cell fibrogenic conversion. Aging Cell 16, 518–528, doi:10.1111/acel.12578 (2017).

29 Ameen, V. & Robson, L. G. Experimental models of duchenne muscular dystrophy: relationship with cardiovascular disease. Open Cardiovasc Med J 4, 265–277, doi:10.2174/1874192401004010265 (2010).

30 Bersini, S. et al. Tackling muscle fibrosis: From molecular mechanisms to next generation engineered models to predict drug delivery. Advanced drug delivery reviews 129, 64–77, doi:10.1016/j.addr.2018.02.009 (2018).

31 Khodadadeh, S., McClelland, M. R., Patrick, J. H., Edwards, R. H. & Evans, G. A. Knee moments in Duchenne muscular dystrophy. Lancet 2, 544–545, doi:10.1016/s0140-6736(86)90114-5 (1986).

32 Muntoni, F., Mateddu, A., Marchei, F., Clerk, A. & Serra, G. Muscular weakness in the mdx mouse. Journal of the neurological sciences 120, 71–77, doi:10.1016/0022-510x(93)90027-v (1993).

33 Ohlendieck, K. et al. Duchenne muscular dystrophy: deficiency of dystrophin-associated proteins in the sarcolemma. Neurology 43, 795–800, doi:10.1212/wnl.43.4.795 (1993).

34 Hicks, M. R. et al. ERBB3 and NGFR mark a distinct skeletal muscle progenitor cell in human development and hPSCs. Nat Cell Biol 20, 46–57, doi:10.1038/s41556-017-0010-2 (2018).

35 Kreitzer, F. R. et al. A robust method to derive functional neural crest cells from human pluripotent stem cells. Am J Stem Cells 2, 119–131 (2013).

36 Sadelain, M., Papapetrou, E. P. & Bushman, F. D. Safe harbours for the integration of new DNA in the human genome. Nat Rev Cancer 12, 51–58, doi:10.1038/nrc3179 (2011).

37 Sweeney, S. M. et al. Candidate cell and matrix interaction domains on the collagen fibril, the predominant protein of vertebrates. The Journal of biological chemistry 283, 21187–21197, doi:10.1074/jbc.M709319200 (2008).

38 Knott, L. & Bailey, A. J. Collagen cross-links in mineralizing tissues: a review of their chemistry, function, and clinical relevance. Bone 22, 181–187 (1998).

39 Philp, C. J. et al. Extracellular Matrix Cross-Linking Enhances Fibroblast Growth and Protects against Matrix Proteolysis in Lung Fibrosis. Am J Respir Cell Mol Biol 58, 594–603, doi:10.1165/rcmb.2016-0379OC (2018).

40 Yoshida, K. et al. Quantitative evaluation of collagen crosslinks and corresponding tensile mechanical properties in mouse cervical tissue during normal pregnancy. PloS one 9, e112391, doi:10.1371/journal.pone.0112391 (2014).

41 Jones, M. G. et al. Nanoscale dysregulation of collagen structure-function disrupts mechano-homeostasis and mediates pulmonary fibrosis. Elife 7, doi:10.7554/eLife.36354 (2018).

42 Eyre, D. R., Paz, M. A. & Gallop, P. M. Cross-linking in collagen and elastin. Annu Rev Biochem 53, 717–748, doi:10.1146/annurev.bi.53.070184.003441 (1984).

43 Deschesnes, R. G., Huot, J., Valerie, K. & Landry, J. Involvement of p38 in apoptosis-associated membrane blebbing and nuclear condensation. Mol Biol Cell 12, 1569–1582, doi:10.1091/mbc.12.6.1569 (2001).

44 Xie, K., Yang, Y. & Jiang, H. Controlling Cellular Volume via Mechanical and Physical Properties of Substrate. Biophys J 114, 675–687, doi:10.1016/j.bpj.2017.11.3785 (2018).

45 Fackler, O. T. & Grosse, R. Cell motility through plasma membrane blebbing. The Journal of cell biology 181, 879–884, doi:10.1083/jcb.200802081 (2008).

46 Colotta, F., Polentarutti, N., Sironi, M. & Mantovani, A. Expression and involvement of c-fos and c-jun protooncogenes in programmed cell death induced by growth factor deprivation in lymphoid cell lines. J Biol Chem 267, 18278–18283 (1992).

47 Zhou, H. et al. Role of c-Fos/JunD in protecting stress-induced cell death. Cell Prolif 40, 431–444, doi:10.1111/j.1365-2184.2007.00444.x (2007).

48 Ross, J. et al. Defects in glycosylation impair satellite stem cell function and niche composition in the muscles of the dystrophic Large(myd) mouse. Stem Cells 30, 2330–2341, doi:10.1002/stem.1197 (2012).

49 Kanagawa, M. et al. Disruption of perlecan binding and matrix assembly by post-translational or genetic disruption of dystroglycan function. FEBS Lett 579, 4792–4796, doi:10.1016/j.febslet.2005.07.059 (2005).

50 P.D., C. H. W. D. A. Y. Laminin polymerization induces a receptor-cytoskeleton network. J Cell Biol. 145, 619–631.

51 Naba, A. et al. The extracellular matrix: Tools and insights for the “omics” era. Matrix biology : journal of the International Society for Matrix Biology 49, 10–24, doi:10.1016/j.matbio.2015.06.003 (2016).

52 Carberry, S., Zweyer, M., Swandulla, D. & Ohlendieck, K. Proteomics reveals drastic increase of extracellular matrix proteins collagen and dermatopontin in the aged mdx diaphragm model of Duchenne muscular dystrophy. International journal of molecular medicine 30, 229–234, doi:10.3892/ijmm.2012.1006 (2012).

53 Holland, A. et al. Comparative Label-Free Mass Spectrometric Analysis of Mildly versus Severely Affected mdx Mouse Skeletal Muscles Identifies Annexin, Lamin, and Vimentin as Universal Dystrophic Markers. Molecules 20, 11317–11344, doi:10.3390/molecules200611317 (2015).

54 Murphy, S. et al. Proteomic profiling of the dystrophin complex and membrane fraction from dystrophic mdx muscle reveals decreases in the cytolinker desmoglein and increases in the extracellular matrix stabilizers biglycan and fibronectin. J Muscle Res Cell Motil 38, 251–268, doi:10.1007/s10974-017-9478-4 (2017).

55 Barrett, A. S. et al. Hydroxylamine Chemical Digestion for Insoluble Extracellular Matrix Characterization. Journal of proteome research 16, 4177–4184, doi:10.1021/acs.jproteome.7b00527 (2017).

56 Crosbie, R. H., Heighway, J., Venzke, D. P., Lee, J. C. & Campbell, K. P. Sarcospan, the 25-kDa transmembrane component of the dystrophin-glycoprotein complex. The Journal of biological chemistry 272, 31221–31224 (1997).

57 Crosbie, R. H. et al. Membrane targeting and stabilization of sarcospan is mediated by the sarcoglycan subcomplex. The Journal of cell biology 145, 153–165 (1999).

58 Marshall, J. L. et al. Dystrophin and utrophin expression require sarcospan: loss of alpha7 integrin exacerbates a newly discovered muscle phenotype in sarcospan-null mice. Human molecular genetics 21, 4378–4393, doi:10.1093/hmg/dds271 (2012).

59 Peter, A. K., Miller, G. & Crosbie, R. H. Disrupted mechanical stability of the dystrophin-glycoprotein complex causes severe muscular dystrophy in sarcospan transgenic mice. Journal of cell science 120, 996–1008, doi:10.1242/jcs.03360 (2007).

60 Gibbs, E. M. et al. High levels of sarcospan are well tolerated and act as a sarcolemmal stabilizer to address skeletal muscle and pulmonary dysfunction in DMD. Hum Mol Genet 25, 5395–5406, doi:10.1093/hmg/ddw356 (2016).

61 Parvatiyar, M. S. et al. Stabilization of the cardiac sarcolemma by sarcospan rescues DMD-associated cardiomyopathy. JCI Insight 5, doi:10.1172/jci.insight.123855 (2019).

62 Parvatiyar, M. S. et al. Sarcospan Regulates Cardiac Isoproterenol Response and Prevents Duchenne Muscular Dystrophy-Associated Cardiomyopathy. J Am Heart Assoc 4, doi:10.1161/JAHA.115.002481 (2015).

63 Chan, J. et al. Galectin-1 induces skeletal muscle differentiation in human fetal mesenchymal stem cells and increases muscle regeneration. Stem Cells 24, 1879–1891, doi:10.1634/stemcells.2005-0564 (2006).

64 Georgiadis, V. et al. Lack of galectin-1 results in defects in myoblast fusion and muscle regeneration. Dev Dyn 236, 1014–1024, doi:10.1002/dvdy.21123 (2007).

65 Van Ry, P. M., Wuebbles, R. D., Key, M. & Burkin, D. J. Galectin-1 Protein Therapy Prevents Pathology and Improves Muscle Function in the mdx Mouse Model of Duchenne Muscular Dystrophy. Molecular therapy : the journal of the American Society of Gene Therapy 23, 1285–1297, doi:10.1038/mt.2015.105 (2015).

66 Watt, D. J., Jones, G. E. & Goldring, K. The involvement of galectin-1 in skeletal muscle determination, differentiation and regeneration. Glycoconj J 19, 615–619, doi:10.1023/B:GLYC.0000014093.23509.92 (2002).

67 Urciuolo, A. et al. Collagen VI regulates satellite cell self-renewal and muscle regeneration. Nature communications 4, 1964, doi:10.1038/ncomms2964 (2013).

68 Sabatelli, P. et al. Expression of collagen VI alpha5 and alpha6 chains in human muscle and in Duchenne muscular dystrophy-related muscle fibrosis. Matrix biology : journal of the International Society for Matrix Biology 31, 187–196, doi:10.1016/j.matbio.2011.12.003 (2012).

69 Hardy, D. et al. Comparative Study of Injury Models for Studying Muscle Regeneration in Mice. PLoS One 11, e0147198, doi:10.1371/journal.pone.0147198 (2016).

70 Campbell, K. P. & Kahl, S. D. Association of dystrophin and an integral membrane glycoprotein. Nature 338, 259–262, doi:10.1038/338259a0 (1989).

71 Ervasti, J. M., Ohlendieck, K., Kahl, S. D., Gaver, M. G. & Campbell, K. P. Deficiency of a glycoprotein component of the dystrophin complex in dystrophic muscle. Nature 345, 315–319, doi:10.1038/345315a0 (1990).

72 Thomas, K., Engler, A. J. & Meyer, G. A. Extracellular matrix regulation in the muscle satellite cell niche. Connective tissue research 56, 1–8, doi:10.3109/03008207.2014.947369 (2015).

73 Engler, A. J., Sen, S., Sweeney, H. L. & Discher, D. E. Matrix elasticity directs stem cell lineage specification. Cell 126, 677–689, doi:10.1016/j.cell.2006.06.044 (2006).

74 Gilbert, P. M. et al. Substrate elasticity regulates skeletal muscle stem cell self-renewal in culture. Science 329, 1078–1081, doi:10.1126/science.1191035 (2010).

75 Engler, A. J. et al. Myotubes differentiate optimally on substrates with tissue-like stiffness: pathological implications for soft or stiff microenvironments. J Cell Biol 166, 877–887, doi:10.1083/jcb.200405004 (2004).

76 Rehfeldt, F., Engler, A. J., Eckhardt, A., Ahmed, F. & Discher, D. E. Cell responses to the mechanochemical microenvironment--implications for regenerative medicine and drug delivery. Adv Drug Deliv Rev 59, 1329–1339, doi:10.1016/j.addr.2007.08.007 (2007).

77 Giordani, L. et al. High-Dimensional Single-Cell Cartography Reveals Novel Skeletal Muscle-Resident Cell Populations. Mol Cell 74, 609–621 e606, doi:10.1016/j.molcel.2019.02.026 (2019).

78 Baghdadi, M. B. et al. Reciprocal signalling by Notch-Collagen V-CALCR retains muscle stem cells in their niche. Nature 557, 714–718, doi:10.1038/s41586-018-0144-9 (2018).

79 Rooney, J. E., Gurpur, P. B. & Burkin, D. J. Laminin-111 protein therapy prevents muscle disease in the mdx mouse model for Duchenne muscular dystrophy. Proceedings of the National Academy of Sciences of the United States of America 106, 7991–7996, doi:10.1073/pnas.0811599106 (2009).

80 Vo, A. H. & McNally, E. M. Modifier genes and their effect on Duchenne muscular dystrophy. Current opinion in neurology 28, 528–534, doi:10.1097/WCO.0000000000000240 (2015).

81 Heydemann, A. et al. Latent TGF-beta-binding protein 4 modifies muscular dystrophy in mice. The Journal of clinical investigation 119, 3703–3712, doi:10.1172/JCI39845 (2009).

82 Capote, J. et al. Osteopontin ablation ameliorates muscular dystrophy by shifting macrophages to a pro-regenerative phenotype. The Journal of cell biology 213, 275–288, doi:10.1083/jcb.201510086 (2016).

83 Afshar Bakooshli, M., et al. A 3D culture model of innervated human skeletal muscle enables studies of the adult neuromuscular junction. Elife 8, doi:10.7554/eLife.44530 (2019).

84 Davoudi, S., et al. MEndR: An In Vitro Functional Assay to Predict In Vivo Muscle Stem Cell-Mediated Repair. Adv Funct Mater 32, doi:ARTN 2106548 10.1002/adfm.202106548 (2022).

85 Marshall, J. L. et al. Sarcospan integration into laminin-binding adhesion complexes that ameliorate muscular dystrophy requires utrophin and alpha7 integrin. Hum Mol Genet 24, 2011–2022, doi:10.1093/hmg/ddu615 (2015).

86 Young, C. S. et al. A Single CRISPR-Cas9 Deletion Strategy that Targets the Majority of DMD Patients Restores Dystrophin Function in hiPSC-Derived Muscle Cells. Cell Stem Cell 18, 533–540, doi:10.1016/j.stem.2016.01.021 (2016).

87 Marshall, J. L. et al. Sarcospan-dependent Akt activation is required for utrophin expression and muscle regeneration. The Journal of cell biology 197, 1009–1027, doi:10.1083/jcb.201110032 (2012).

88 van Zwieten, R. W. et al. Assessing dystrophies and other muscle diseases at the nanometer scale by atomic force microscopy. Nanomedicine 9, 393–406, doi:10.2217/NNM.12.215 (2014).

89 Sharma, S. et al. Nanoscale characterization of effect of L-arginine on Streptococcus mutans biofilm adhesion by atomic force microscopy. Microbiology 160, 1466–1473, doi:10.1099/mic.0.075267-0 (2014).

90 Dobin, A. et al. STAR: ultrafast universal RNA-seq aligner. Bioinformatics 29, 15–21, doi:10.1093/bioinformatics/bts635 (2013).

91 Robinson, M. D., McCarthy, D. J. & Smyth, G. K. edgeR: a Bioconductor package for differential expression analysis of digital gene expression data. Bioinformatics 26, 139–140, doi:10.1093/bioinformatics/btp616 (2010).

92 Wither, M. J., Hansen, K. C. & Reisz, J. A. Mass Spectrometry-Based Bottom-Up Proteomics: Sample Preparation, LC-MS/MS Analysis, and Database Query Strategies. Curr Protoc Protein Sci 86, 16 14 11–16 14 20, doi:10.1002/cpps.18 (2016).

93 Johnson, T. D. et al. Quantification of decellularized human myocardial matrix: A comparison of six patients. Proteomics Clin Appl 10, 75–83, doi:10.1002/prca.201500048 (2016).

94 Xia, J., Sinelnikov, I. V., Han, B. & Wishart, D. S. MetaboAnalyst 3.0--making metabolomics more meaningful. Nucleic Acids Res 43, W251–257, doi:10.1093/nar/gkv380 (2015).

