## Supplemental Figures for "Myoscaffolds reveal laminin scarring is detrimental for stem cell function while sarcospan induces compensatory fibrosis"

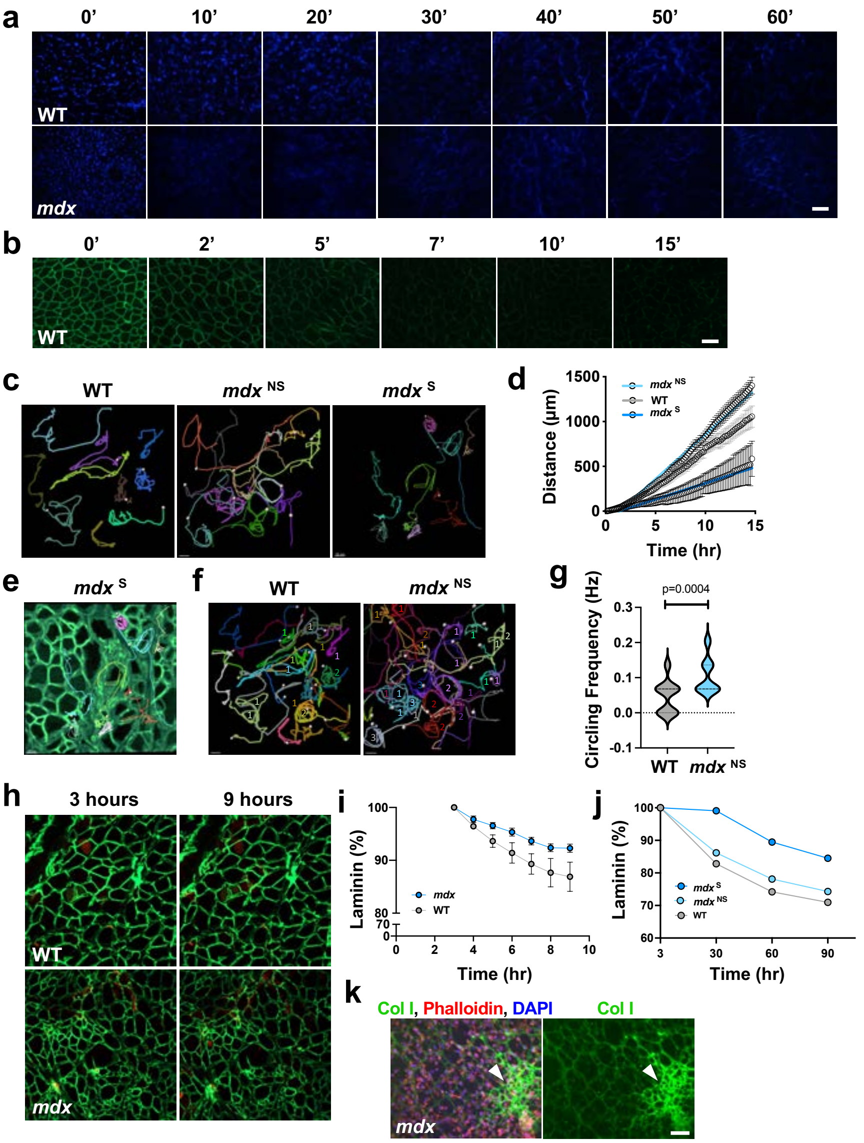


**Supplemental Figure 1. Myoscaffolds from healthy and diseased muscle differentially support SMPC migration and growth.**
**a.** Representative images of DAPI staining (blue) performed on transverse cryosections from wild-type (WT) and *mdx* quadriceps muscles that were decellularized with 1% SDS solution for the indicated times (10’ to 60’). Non-decellularized (whole) muscle sections were used as controls and are denoted as the 0’ time point. Scale bar, 100 µm. (n=3 independent experiments) **b.** Immunofluorescence analysis of dystrophin (green) shows rapid removal of the protein within 15 minutes of 1% SDS treatment (indicated times 2’ to 15’). Non-decellularized (whole) muscle sections were used as controls (0’) (n=3 independent experiments). Scale bar, 100 µm. **c.** Individual SMPC tracks of total displacement are marked by unique colors and highlight SMPC behavior differences on *mdx* myoscaffolds (WTC-11 cells). **d.** Total cell displacement (microns) was calculated by combining cell speed with cumulative sum of speeds from previous time points. Non-constrained linear regression was used to calculate the slope of displacement (WTC-11 cells, n=10-14 cells/tissue; based on observations from n=3 independent experiments) **e.** Overlay of the cell tracks from panel (**c**) with the image of the *mdx* $^{S}$ region from live cell imaging (Video S3) showing limited cell motility over fibrotic scars. **f.** The frequency at which SMPCs circled the basement membrane was calculated for individual SMPCs on WT and *mdx* $^{NS}$ myoscaffolds (WTC-11 cells, n=19-22 cells/tissue; based on observations from n=3 independent experiments). The white dots represent the final location of the cell following the 15-hour tracking period. **g.** Circling frequency was calculated as the number of times a cell circled the basement membrane over a 14 hour period. Differences in frequency were calculated using an independent samples t-test ($p$<0.05). We found that cells on *mdx* $^{NS}$ myoscaffolds circled the basement membrane at a 2.3 fold greater frequency than those on WT myoscaffolds (WT: 0.047 ± 0.046 vs. *mdx* $^{NS}$: 0.105 ± 0.050 Hz (mean ± st. dev.), $p$=0.0004) (WTC-11 cells, n=19-22 cells/tissue). **h.** Representative immunofluorescent images of WT and *mdx* myoscaffolds cultured with SMPCs (WTC-11 hiPSCs) taken at 3 and 9 hours after cell seeding (laminin alpha 2-green, SMPCs- red). **i.** Mean laminin intensity was calculated each hour over the entire field of view, starting after 3 hours, which was the time required for the image to stabilize as cells settled and migrated on the myoscaffolds. Although not significant, there was a trend toward a greater decrease in laminin intensity in WT myoscaffolds, compared to *mdx*, indicating greater laminin degradation (n=4 samples/group, mean +/- SEM). **j.** For a single experiment, laminin fluorescence intensity was quantified over 90 hours of imaging, starting 3 hours after cell seeding. Laminin intensity began to drop immediately on WT and *mdx* $^{NS}$ myoscaffolds, suggestive of laminin degradation, while laminin intensity dropped only 1% in the first 30 hours when cells were cultured on *mdx* $^{S}$ regions (n=1 region/tissue type, based on observations from n=3 independent experiments). **i.** Representative image of an *mdx* myoscaffold with a fibrotic scar (white arrowhead), stained for collagen I (Col I, green), phalloidin (red), and DAPI (blue). Collagen I in the fibrotic scar was resistant to remodeling by SMPCs, as indicated by increased fluorescence in the fibrotic region relative to the rest of the myoscaffolds (n=4 independent experiments). Scale bar, 100 µm.

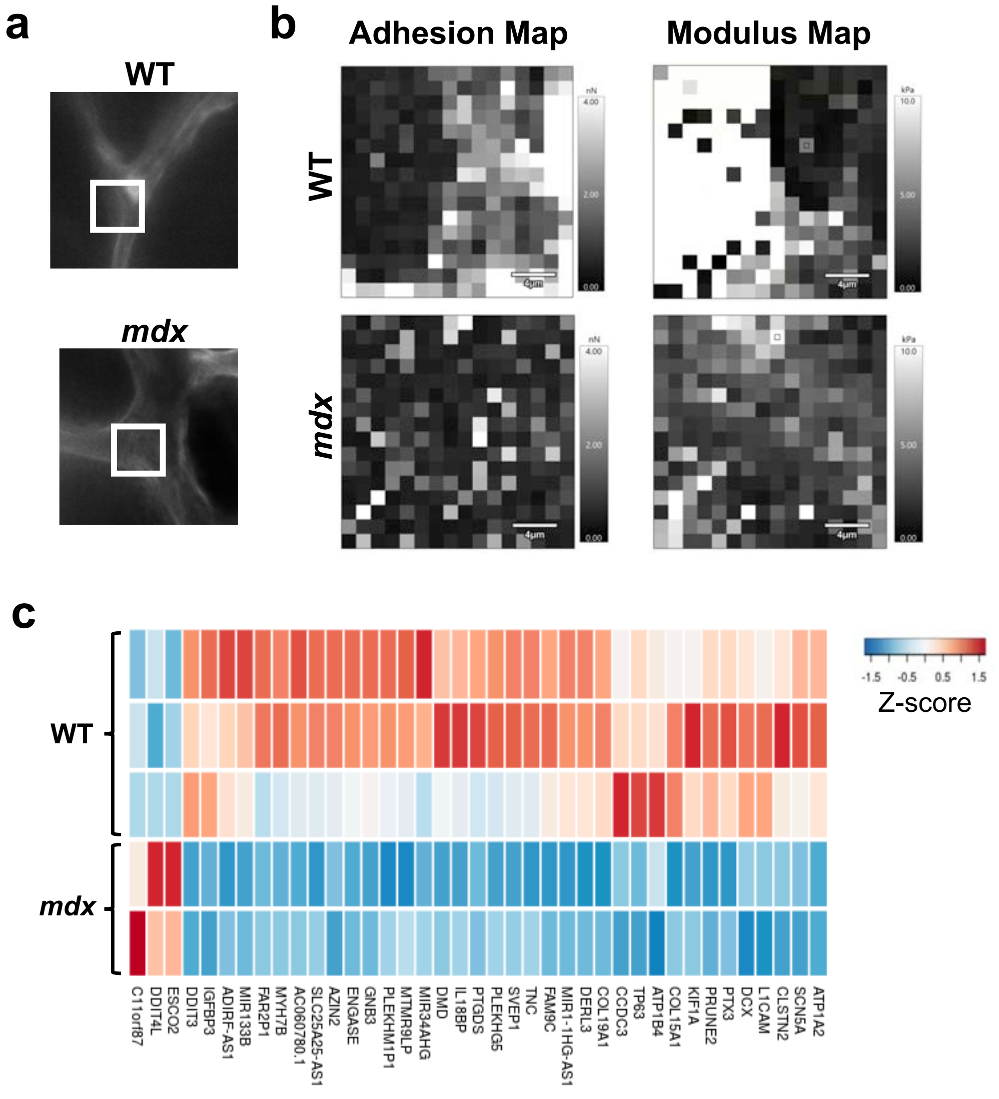


**Supplemental Figure 2. Stiff *mdx* myoscaffolds have reduced adhesive properties and induce downregulation of genes associated with cell adhesion, migration, and skeletal muscle maturation.**

**a.** Images of the WT and *mdx* ECM samples from AFM testing stained for laminin using indirect immunofluorescence. The white box represents the 20 x 20µm region scanned from each sample. **b.** Representative adhesion and modulus maps from the samples shown in (**a**) (n=3 independent experiments). **c.** RNA sequencing was performed using total RNA prepared from SMPCs cultured (5 days) on WT and *mdx* myoscaffolds. Expression patterns of differentially expressed genes (DEGs) from SMPCs (H9 cells) cultured on WT (n=3) and *mdx* (n=2) myoscaffolds. Expression levels are shown as Z-scores, as in scale. Most DEGs were downregulated in *mdx* compared to WT, including L1 cell adhesion molecule (L1CAM) and calsyntenin (CLSTN2) that play roles in cell adhesion and migration. Genes associated with skeletal muscle development and maturation such as myosin heavy chain 7B (*MYH7B*), dystrophin (*DMD*), tenacin C (*TNC*), and kinesin family member 1A (*KIF1A*) were also downregulated. Several cell adhesion genes including integrins (*ITGA4, ITGA6*), M-cadherin (*CDH15*), and neural cell adhesion marker (*NCAM1*) as well as sarcomeric genes (myosin heavy chain 3 and 7 (*MYH3, MYH7*) and troponin T2 (*TNNT2*) trended toward downregulation in SMPCs cultured on *mdx* myoscaffolds ($p<0.05$). Genes associated with basement membrane assembly, including collagens type XV and XIX (COL15A1 and COL19A1), exhibited decreased expression in *mdx* samples.

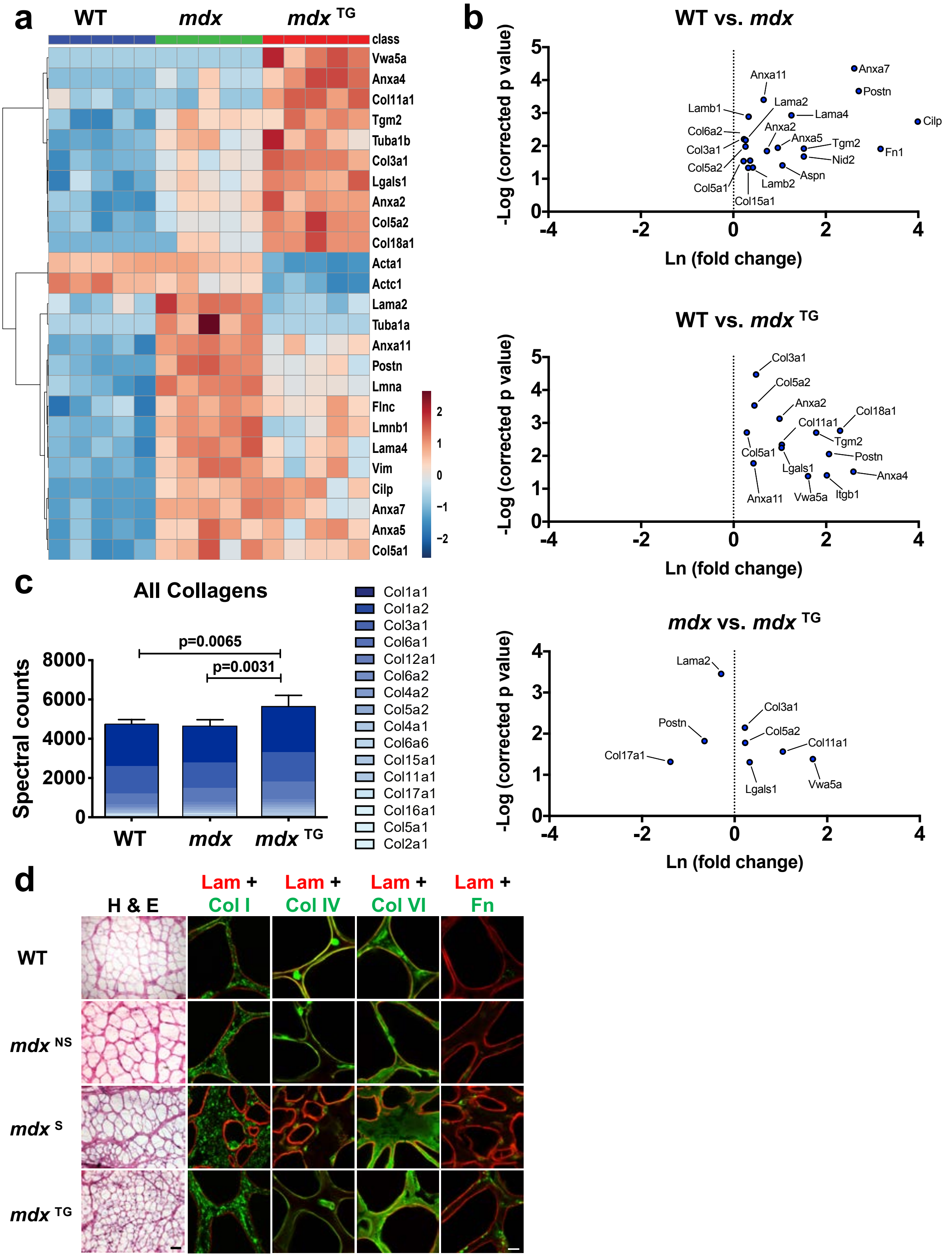


**Supplemental Figure 3. The matrisome of *mdx* and *mdx* $^{TG}$ samples diverge from that of wild-type muscle.**
**a.** Heat map from ECM focused proteomics reveals distinct clustering of each phenotype (n=5 samples/group). **b.** Volcano plots showing ln fold change plotted against −log10 adjusted P value for WT vs. *mdx,* WT vs. *mdx* $^{TG}$, and *mdx* vs. *mdx* $^{TG}$ samples. **c.** Column graphs showing the abundance of collagens in skeletal muscle from WT, *mdx*, and *mdx* $^{TG}$ samples. *P* values reflect analysis by one-way ANOVA. **d.** Replicate images from analysis of the abundance and organization of ECM components in the basement membrane and interstitial matrix regions of WT, *mdx*, and *mdx* $^{TG}$ samples. Regions of *mdx* myoscaffolds without fibrotic scars (*mdx* $^{NS}$) and with scars (*mdx* $^{S}$) shown separately. Representative images are shown from H&E staining, along with indirect fluorescent image analysis of myoscaffolds co-stained for laminin α2 (Lam) (red) with collagen I (Col I), IV (Col IV), VI (Col VI), and fibronectin (Fn) (green), respectively (selected images from n=4 independent experiments). Scale bars, 100μm (H&E) and 8μm (IFA).

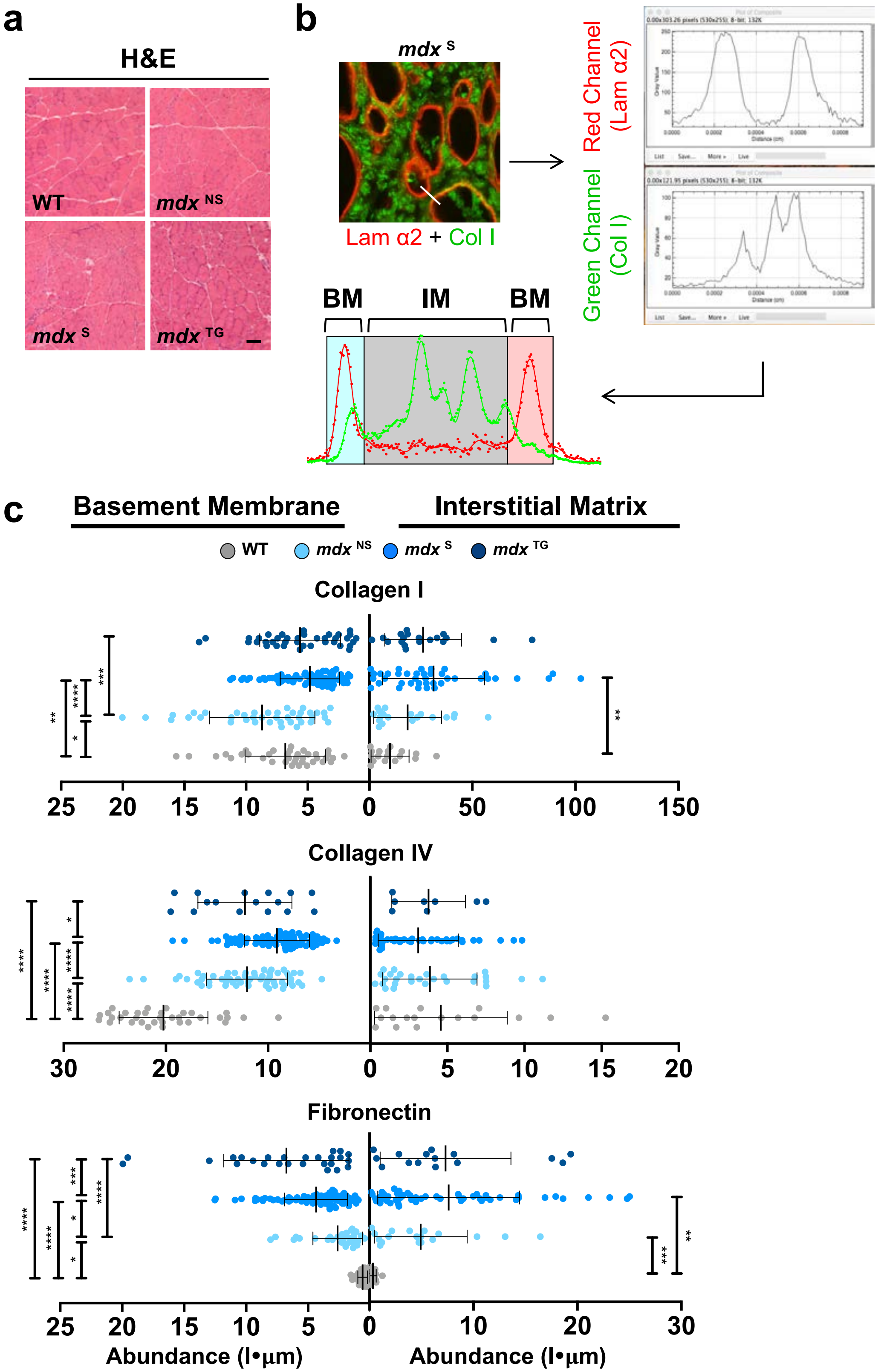


**Supplemental Figure 4. Quantification of ECM protein abundance and distribution reveals alterations in healthy and diseased tissues.** **a.** Representative images of transverse sections of the quadriceps muscle from WT, *mdx* and *mdx* TG mice stained with H&E to visualize muscle pathology. **b.** To quantify the distribution of ECM proteins in the basement membrane (BM) and interstitial matrix (IM) of co-stained myoscaffolds, a line was drawn across the endomysium (white bar) on confocal images and the line scan function in Image J was used to generate plot profiles of the pixel intensity for each ECM protein (sample confocal image of an *mdx* S myoscaffold: red channel-laminin α2, green channel-collagen I). The plot profiles were exported into Matlab and a custom algorithm was utilized to calculate the abundance of each protein in the BM and IM regions. **c.** Graphs showing the abundance of collagens I and IV and fibronectin in the BM and IM of WT, *mdx* NS, *mdx* S, and *mdx* TG ECM (n=20-40 measurements/group for each protein, n=4 independent experiments). Between group differences were analyzed by one-way ANOVA. *P* values are as follows: * = *p<0.05*, ** = *p<0.01*, *** = *p<0.001*, **** = *p<0.0001*.

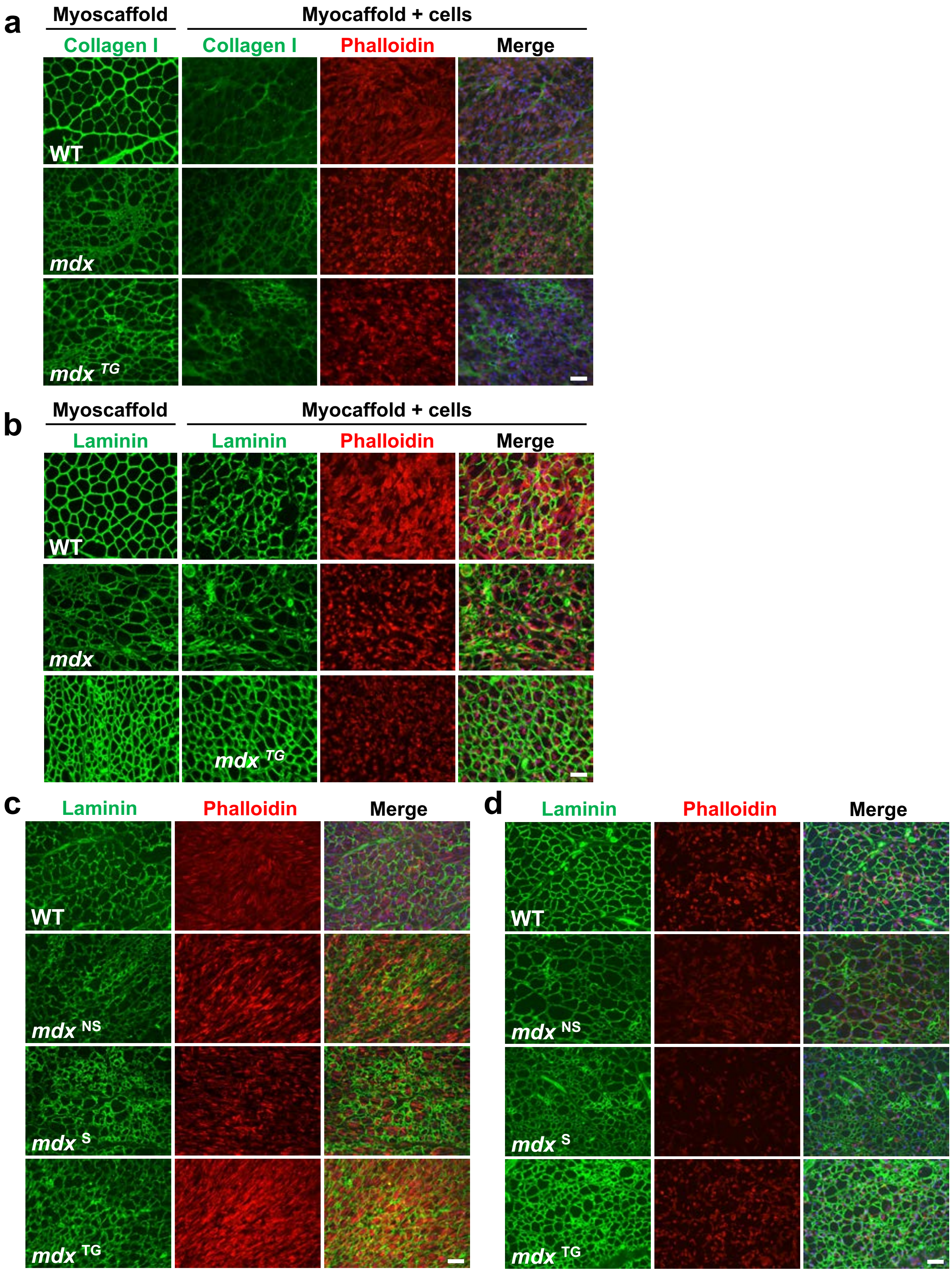


**Supplemental Figure 5. SMPCs do not remodel laminin in *mdx* fibrotic scars.**
**a-b.** Indirect immunofluorescence confocal microscopy of SMPCs (CDMD 1002) cultured for 5 days on WT, *mdx,* and *mdx* $^{TG}$ myoscaffolds (myoscaffold + cells) stained with antibodies recognizing collagen I (**a**) or laminin (**b-d**) (green), along with phalloidin (red) and DAPI (blue). The fluorescence intensity was only controlled between the myoscaffolds and myoscaffolds + cells condition for each group (WT, *mdx,* and *mdx* $^{TG}$), therefore laminin and collagen I abundance cannot be compared between groups. Scale bar, 100µm. **c-d.** Immunofluorescent images from replicate experiments performed with SMPCs (H9) and stained as above. The same muscle samples were used to generate the myoscaffolds shown in panels (**c**) and (**d**), but 2 different clones of the H9 cell line were used (clone 1 in panel **c**, clone 2 in panel **d**). Although the two clones exhibit variable cell proliferation and laminin remodeling capacity, they demonstrate similar trends in laminin remodeling and cell morphology, including reduced laminin remodeling on *mdx* $^{S}$ myoscaffolds.

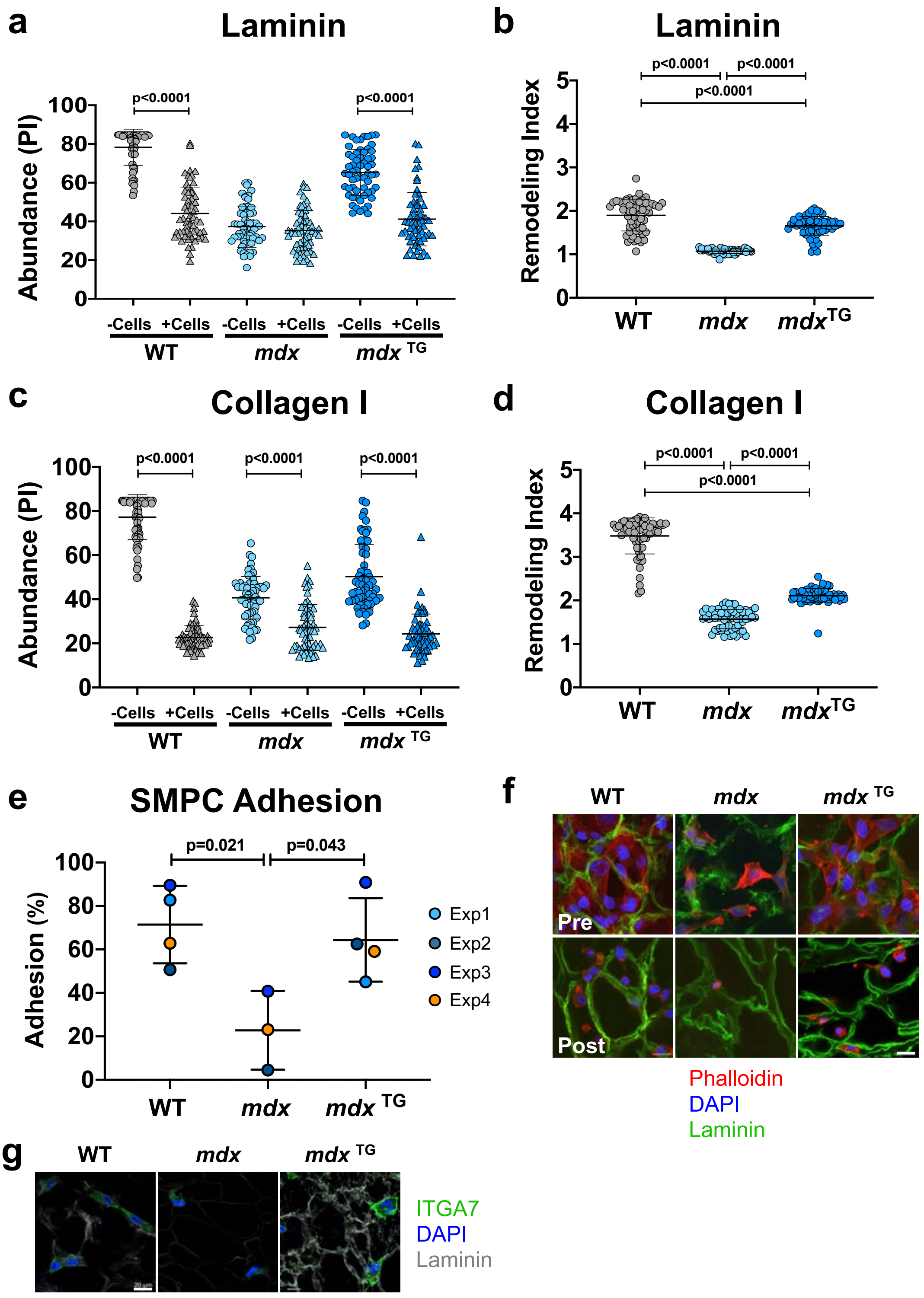


**Supplemental Figure 6. Improved SMPC adhesion to *mdx* TG myoscaffolds accompanies improved laminin remodeling and is facilitated by increased expression of Itga7.**

**a-d.** WT, *mdx*, and *mdx* TG myoscaffolds without (- cells) and with SMPCs (+ cells, CDMD 1002 cells) were cultured in proliferation media (5 days) and then stained with antibodies against laminin and collagen I. Protein abundance (maximum pixel intensity (PI)) was measured from 60 endomysial locations/ myoscaffold. The remodeling index (RI) was calculated as the ratio of protein abundance (PI) in the myoscaffolds in the absence of cells to protein abundance in the myoscaffolds after cell seeding. SMPCs were unable to remodel laminin in *mdx* scaffolds (**a**), with an RI value close to 1 (**b**). While all SMPCs remodeled collagen I (**c**), the RI was significantly lower for cells on *mdx* myoscaffolds (**d**). **e.** Average SMPC adhesion to WT, *mdx*, and *mdx* TG myoscaffolds was calculated from n=4 independent experiments using both CDMD 1002 (Exp1) and H9 cell lines (Exp2, Exp3, and Exp4). A significant reduction in cell adhesion was observed with culture on *mdx* scaffolds, while there was no difference observed between WT and *mdx* TG across all cell lines. **f.** Images of SMPCs (H9 cells) cultured for 4 hours on WT, *mdx*, and *mdx* TG myoscaffolds both before (pre) and after (post) exposure to dissociation buffer. Scale bar, 20 µm. **g.** An increased abundance of integrin α7 (Itga7) was observed in SMPCs (H9 cells) cultured for 4 hours on *mdx* TG myoscaffolds, compared to those on *mdx* scaffolds. Scale bar, 20 µm. (n=3 independent experiments)

a

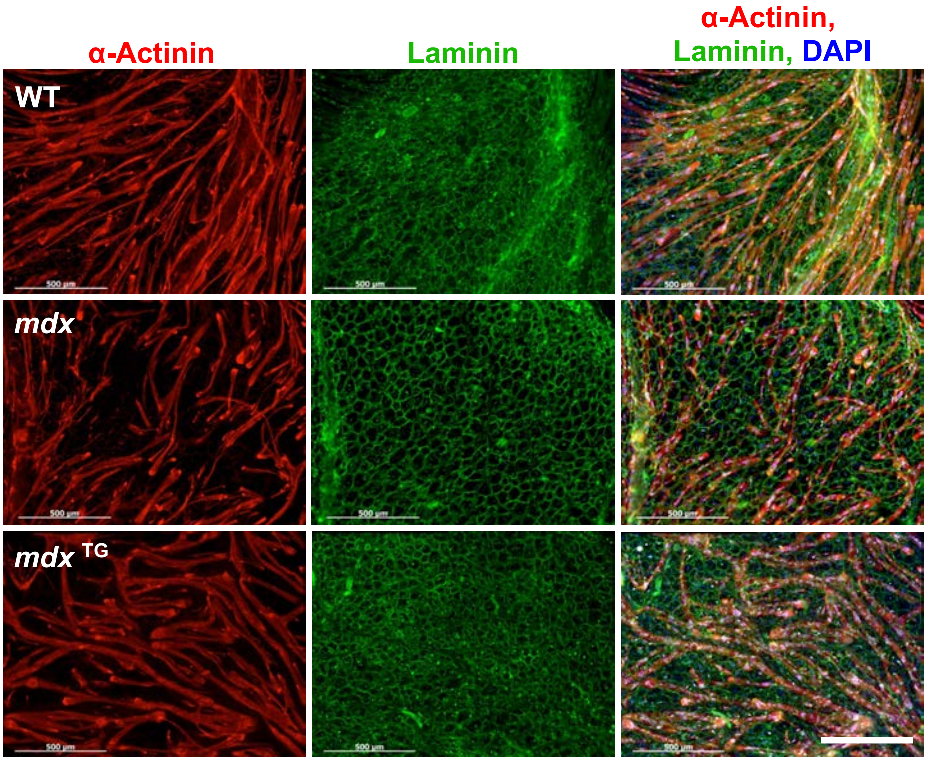


b

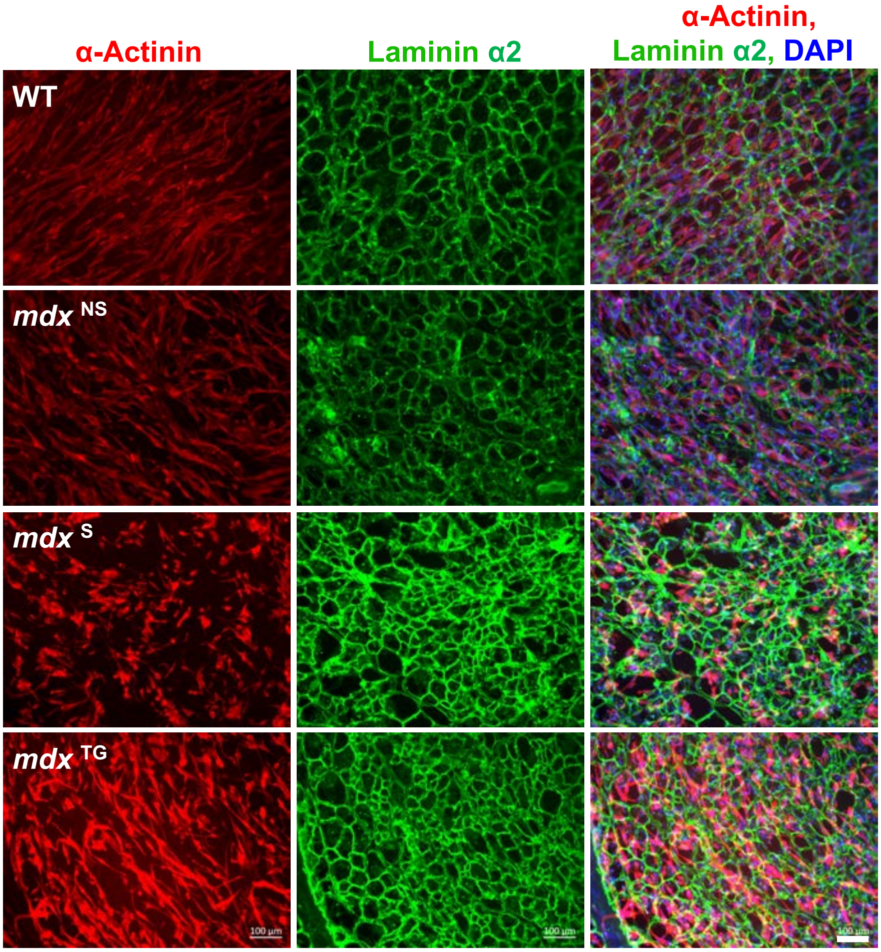


**Supplemental Figure 7. SMPC differentiation is inhibited on *mdx* myoscaffolds.**
**a-b**. Replicate SMPC differentiation experiments performed using CDMD 1002 (**a**) and H9 (**b**) cell lines. Cell differentiation is inhibited on *mdx* $^{S}$ myoscaffolds, while both cell lines demonstrate robust myotube formation on *mdx* $^{TG}$ myoscaffolds. Scale bar, 500µm (**a**) and 100µm (**b**).

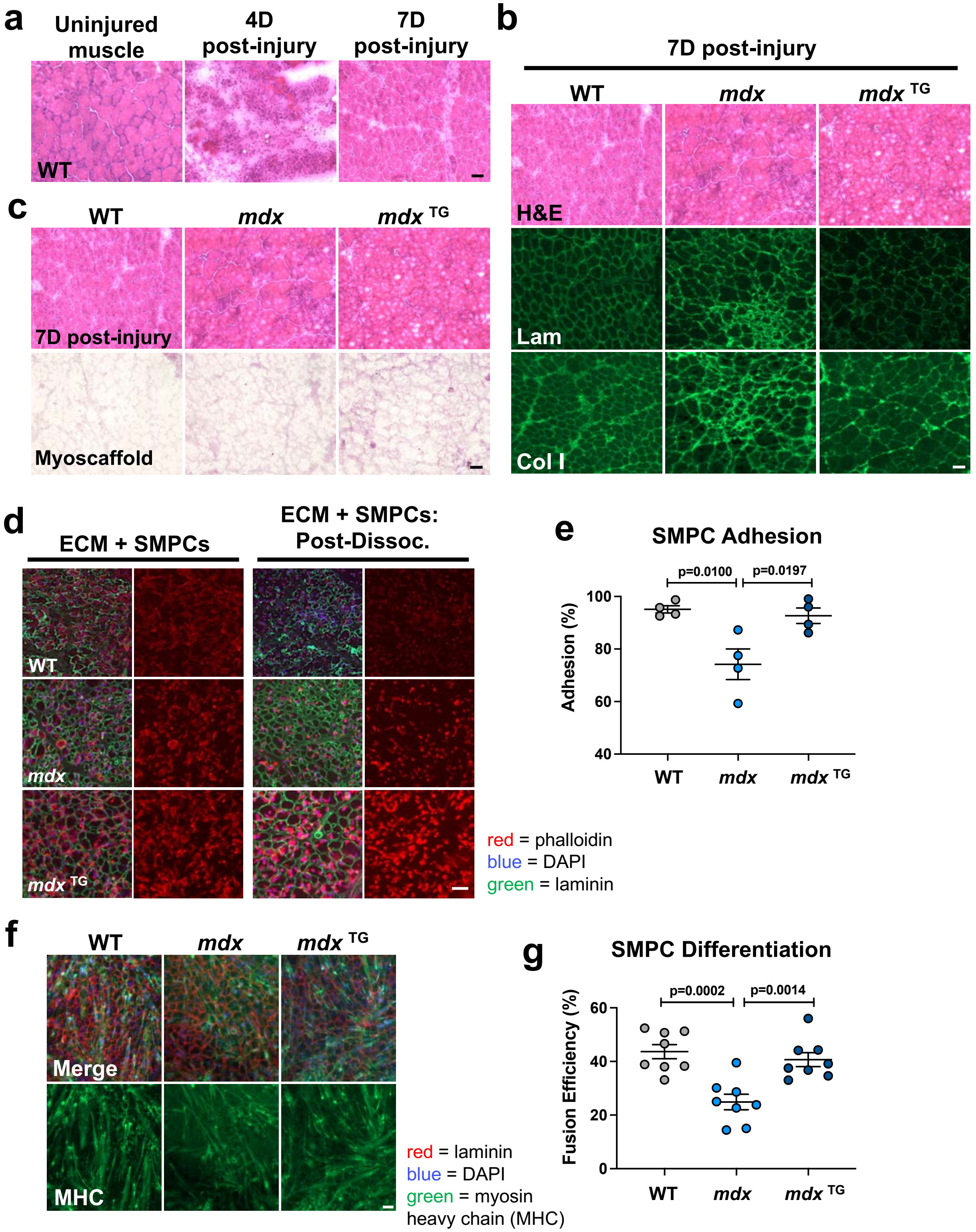


**Supplemental Figure 8. Reduced SMPC adhesion and fusion efficiency on *mdx* myoscaffolds derived from $BaCl_2$ treated muscle.** **a.** Images of H&E stained transverse muscle sections from the tibialis anterior (TA) of wild-type (WT) mice; uninjured and following $BaCl_2$ injection (4 and 7 days post-injury). Scale bar=50µm. **b.** H&E and immunofluorescent images of TA muscle sections from WT, *mdx*, and *mdx* $^{TG}$, taken at 7 days post-injury. Scale bar=50µm. **c.** H&E images showing whole muscle (7D post-injury) and a muscle section decellularized in 1% SDS for 10 minutes (Decell), from WT, *mdx*, and *mdx* $^{TG}$ TA muscles taken at 7 days post-injury. Cellular material is effectively removed after 10 minutes while ECM architecture is maintained. Scale bar=50µm. **d.** Images of SMPCs (H9 cells) cultured for 4 hours on WT, *mdx*, and *mdx* $^{TG}$ myoscaffolds both without (ECM + SMPCs) and with (ECM + SMPCs: Post-Dissoc) exposure to dissociation buffer (n=4/group, H9 SMPCs). **e.** A significant reduction in cell adhesion was observed on *mdx* myoscaffolds relative to both WT and *mdx* $^{TG}$ (n=4/group). Scale bar=100µm. **f.** Immunofluorescent images of SMPCs cultured in proliferation media and then differentiated on WT, *mdx*, and *mdx* $^{TG}$ myoscaffolds (n=4/group, H9 SMPCs). Scale bar=50µm. **g.** Fusion efficiency was significantly reduced on *mdx* myoscaffolds (fusion efficiency calculated from n=4/group, 2 locations per tissue).

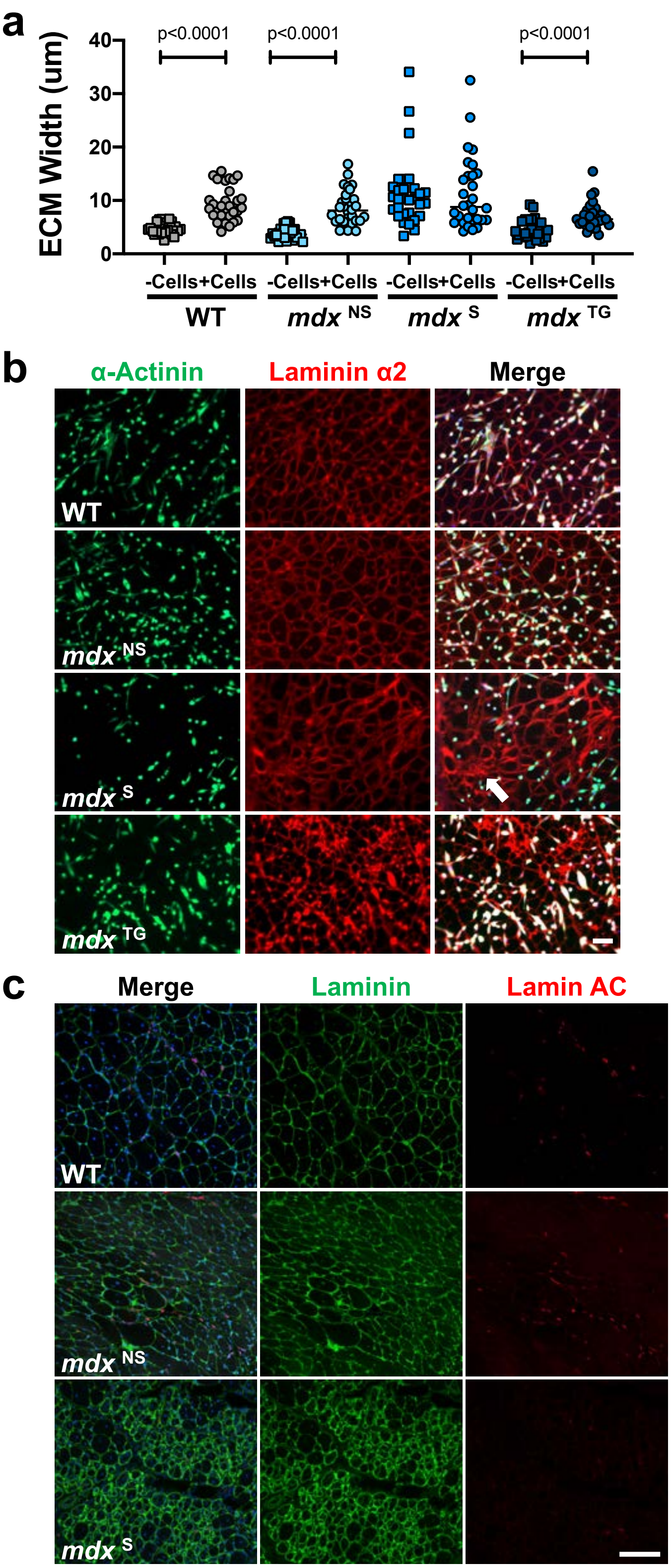


**Supplemental Figure 9. Quantification of endomysial width reveals limited satellite-cell mediated ECM remodeling of *mdx* $^{S}$ myoscaffolds while limited SMPC engraftment is observed in *mdx* $^{S}$ *in vivo*.**

**a.** Confocal images of murine satellite cells differentiated on WT, *mdx* $^{NS}$, *mdx* $^{S}$, and *mdx* $^{TG}$ myoscaffolds (+Cells), as well as control myoscaffolds without cells (-Cells) were utilized to measure endomysial width from 20 locations/image (line drawing tool, Image J) (n=3 samples/group). Myoscaffolds were labeled with antibodies against laminin to demarcate the basement membrane. There was a significant increase in the width of the endomysium in all tissues following cell differentiation, except the *mdx* $^{S}$ myoscaffolds. **b.** Confocal images from replicate cell differentiation experiments utilizing ZsGreen fluorescent murine satellite cells cultured on WT, *mdx* $^{NS}$, *mdx* $^{S}$, and *mdx* $^{TG}$ myoscaffolds and stained with laminin α2 (red). Reduced numbers of SMPCs in laminin dense regions of fibrotic scars (arrow) are observed on *mdx* $^{S}$ scaffolds. Scale bar, 100µm. **c.** Replicate images from SMPC injected C57-NSG and *mdx*-NSG mice, stained for laminin (green), lamin AC (red), and DAPI (blue). SMPCs in WT and regions of the *mdx* muscle without fibrotic scars (*mdx* $^{NS}$) integrated throughout the tissue, while cells were unable to penetrate thickened laminin in fibrotic scars (*mdx* $^{S}$). Scale bar, 100µm.
